# Revisiting oxygen toxicity: evolution and adaptation to superoxide in a SOD-deficient bacterial pathogen

**DOI:** 10.1101/2024.09.25.614947

**Authors:** Samuel G. Huete, Alejandro Leyva, Etienne Kornobis, Thomas Cokelaer, Pierre Lechat, Marc Monot, Rosario Duran, Mathieu Picardeau, Nadia Benaroudj

**Author notes:** Contributing authors.

## Abstract

Defenses against oxidants are crucial for the virulence of pathogens, with superoxide scavenging enzymes (SOSEs) playing a vital role for most aerobes. However, our knowledge of superoxide adaptation primarily stems from the study of SOSE-encoding bacteria. Here, we investigated the evolution of a naturally SOSE-deficient pathogen (*Leptospira* spp.), along with the alternative mechanisms it recruits to combat superoxide stress. We demonstrate that emergence of pathogenic *Leptospira* correlated with SOD loss, but that a long-lasting adaptation to superoxide remains possible. We reveal that cysteine and leucine biosynthesis are the most induced pathways in response to superoxide and demonstrate the importance of sulfur metabolism in superoxide adaptation in this SOSE-deficient model. We also propose cysteine oxidation as a key mediator of superoxide toxicity in the absence of SOSEs. This study challenges our conventional understanding of the oxygen toxicity theory and proposes a new model of superoxide adaptation through metabolic rewiring in bacteria.

## Introduction

Superoxide anion (O_2_^•-^) is a bactericidal oxidant resulting from the one electron-reduction of dioxygen (O_2_) in living organisms. Aerobic pathogens not only encounter the superoxide generated during their own oxygen metabolism, but also high concentrations of host-produced superoxide during infection. Superoxide is spontaneously converted into hydrogen peroxide (H_2_O_2_), but this reaction can be catalyzed by superoxide-scavenging enzymes (SOSEs) like superoxide dismutases (SODs) or reductases (SORs) 10000 times faster^1^. SOSEs are thus considered as the first line of defense against the toxic effects of superoxide, allowing its rapid elimination from the bacterium.

The oxygen toxicity theory, formulated more than 4 decades ago, posits that (i) superoxide is a toxic molecule produced within aerobic organisms and (ii) SODs (and SOSEs in general) are vital for aerobic organisms to detoxify O_2_^•-2,3^. However, this theory was established studying model bacteria and has not been extensively examined across various microorganisms, overlooking SOSE-deficient pathogenic aerobes. How these aerobic bacteria protect themselves against superoxide remains to be elucidated.

The *Leptospira* genus includes both free-living saprophytes as well as pathogenic species responsible for leptospirosis, a re-emerging yet often neglected zoonotic disease^4^. Leptospires are aerobes exposed to endogenous ROS, but only pathogenic species can survive the oxidative stress imposed by the host innate immunity^5^. Specifically, pathogenic *Leptospira* possess a catalase that efficiently degrades H_2_O_2_ and is required for *Leptospira* virulence^6,7^. Previous genomic comparisons within the *Leptospira* genus suggested that certain pathogenic species may lack any SOD-encoding gene^8^, but a large-scale comprehensive analysis was required to confirm this finding.

Here, we investigate the universality of the oxygen toxicity theory to all human bacterial pathogens. We present evidence challenging its assumptions, by showing that two aerobic pathogens (*Leptospira* spp. and *Mycoplasmataceae* spp.) lack conventional superoxide detoxification systems. We also deciphered the evolutionary events behind the loss of SOD in pathogenic *Leptospira* spp. Using a multi-omic approach, we demonstrated that superoxide adaptation remains possible in the absence of SOSEs and characterized the role of cysteine and leucine biosynthesis as the most induced pathways in response to superoxide. Altogether, our findings expand our understanding of how aerobic pathogens respond to oxidative stress, encouraging to revisit the assumptions of the oxygen toxicity theory.

## Results

### Distribution of SOSE among bacteria

To gain a comprehensive understanding of the prevalence of SOSEs in pathogenic bacteria, we analyzed a database of 1110 established bacterial pathogens infecting humans^9^. We identified that over 95% (*N*=1057) of bacterial pathogens encode a SOSE, whereas less than 5% (*N*=53) lack any SOSE in their genome (Fig. 1A, Table S1). Surprisingly, we found a nearly equal distribution between aerobes and anaerobes among SOSE-deficient species. Most aerobic SOSE-deficient pathogens are in the *Mycoplasmataceae* family and the *Leptospira* genus (Fig. 1A). *Mycoplasmataceae* spp. have reduced genomes (0.58-1.73 Mb) and metabolism, including the absence of oxidative phosphorylation^10^. However, *Leptospira* spp., with a complete respiration chain capable of generating superoxide^11^, offer a suitable model for investigating adaptation to superoxide in SOSE-deficient aerobic pathogens.

**Figure 1.**
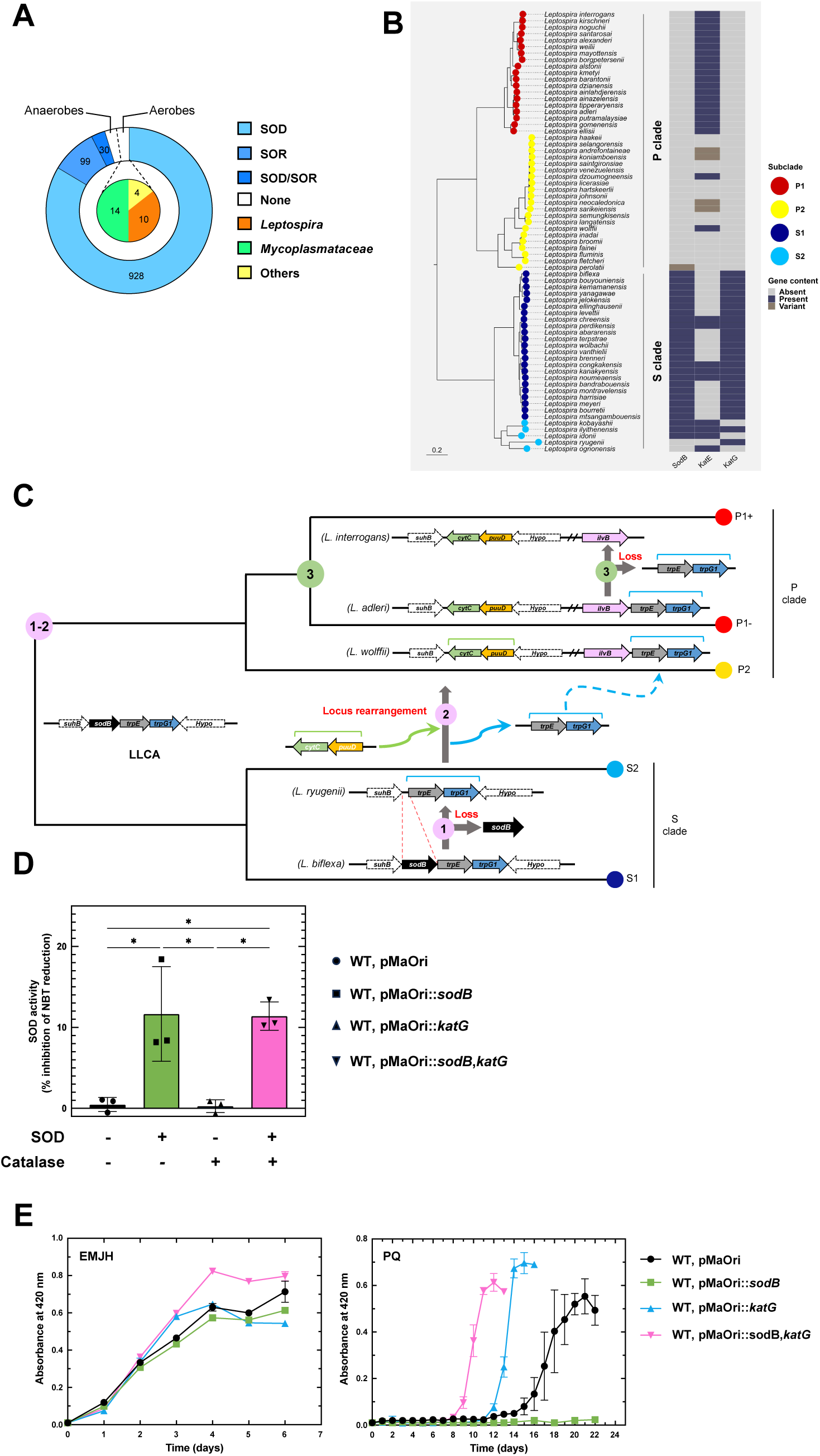
Pathogenic *Leptospira* spp. have lost their SOD. (A) Distribution of SOSEs (SOR or SOD) in different bacterial pathogenic species. SOD and SOR orthologs were searched in a pathogen genome database using a e-value cutoff of 0.01. Aerobicity was indicated for the species that do not contain any SOSE. Numbers correspond to respective number of genomes. (B) Phylogenetic distribution of SOD and catalases (KatE or KatG) in 68 *Leptospira* spp. with an e-value cutoff of 0.01. P1 and P2 species (P clade) are in red and yellow, respectively. S1 and S2 species (S clade) are in dark blue and cyan, respectively. Blue: present, light grey: absent, dark grey: variant. (C) Schematic evolutionary trajectories leading to the loss of SOD in pathogenic *Leptospira* spp. The *sodB* loci are represented in the Last Leptospiral Common Ancestor (LLCA), in S1 (dark blue), S2 (cyan), P2 (yellow), P1-(low virulent) and P1+ (high virulent) (red) species. The three key steps are indicated. (D) SOD activity in the WT (circle), *sodB*-expressing (square), *katG*-expressing (triangle) and *sodB*- and *katG*-expressing (inverted triangle) *L. interrogans* strains. The activity is expressed as the percentage of NBT reduction inhibition per μg of total extract. Data are mean and SD of three independent experiments. *, p-value of 0.01. (E) Growth curve of WT (black circle), *sodB*-expressing (green square), *katG*-expressing (blue triangle) and *sodB*- and *katG*-expressing (pink inverted triangle) *L. interrogans* strains in the presence of 2.5 μM paraquat. Growth was assessed by measure of absorbance at 420 nm. Data are mean and SD of three independent experiments.

### Pathogenic *Leptospira* lost the *sod* gene in the evolution towards pathogenicity

To decipher the evolutionary trajectories leading to the appearance of SOSE-deficient species in *Leptospira*, we investigated the evolution of the SOD and SOR-encoding genes in the *Leptospira* genus and the *Spirochaetes* phylum. This phylum comprises several pathogens in addition to *Leptospira*, such as the causative agents of Lyme disease, relapsing fever and syphilis (*Borrelia burgdorferi*, *Borrelia hermsii*, and *Treponema pallidum*, respectively). We constructed a core-genome phylogeny of the 164 culturable species of the *Spirochaetes* phylum and mapped the presence of a SOD or SOR-encoding ORF (Fig. S1, Table S2). Our findings revealed that 72% (*N*=118) of *Spirochaetes* encode a SOSE. The presence of a SOR does not consistently correlate with the absence of a SOD but it does with anaerobicity (66% of anaerobes have a SOR) (Fig. S1). Among the 28% (*N*=46) of *Spirochaetes* lacking any SOSEs, 89% (*N*=41) are *Leptospira* spp., the remaining 5 being strict anaerobes (Fig. S1). Hence, pathogenic *Leptospira* spp. are the only aerobic *Spirochaetes* lacking a SOSE-encoding gene.

Intriguingly, while most *Leptospira* spp. from the P clade, containing pathogenic species, lack any protein with homology to SOD, a *sod* gene is present in almost all species from the S clade, containing strictly environmental saprophytic species (Fig. 1B, Fig. S2). Evolutionary analysis revealed that the Last Leptospiral Common Ancestor (LLCA) possessed a *sodB* gene, which was maintained in S clades and subsequently lost when *Leptospira* transitioned from S to P clades (Fig. 1C, Fig. S1, Fig. S2). Consistently, all *sodB* genes in *Leptospira* spp. exhibit a highly conserved genetic locus, while the *sodA* homolog in *L. perolatii* was very likely acquired independently through horizontal gene transfer (Fig. S2). Genus-wide comparative genomic analysis of the *sodB*-encoding locus revealed that *trpE* and *trpG1*, the two genes downstream *sodB*, were displaced downstream *ilvB* and substituted with *puuD* and *cytC*. This locus rearrangement probably occurred subsequently to the loss of *sodB* as it is possible to identify S2 species (such as *L. ryugenii*) where *trpE* and *trpG1* are present in their original loci despite of the absence of a *sodB* (Fig. 1C, Fig. S2). The *sodB* loss and *trpE* and *trpG1* displacement likely occurred before the emergence of the P clade (steps 1-2 in Fig. 1C, Fig. S2). Finally, *trpE* and *trpG1* were lost by deletion upon the transition of P1-group (species with low virulence) to the P1+ group (highly virulent species) (step 3 in Fig. 1C, Fig. S2). Altogether, these findings not only establish that *Leptospira* is one of the few SOSE-deficient aerobic pathogens, but also reveal that the loss of its SOD coincided with the transition from saprophytes to host-adapted species.

### Heterologous expression of *sodB* in *L. interrogans* impairs fitness in the presence of superoxide

Given that pathogenic *Leptospira* have lost *sodB*, we tested whether the presence of a SOD activity would improve their tolerance to superoxide. The *sodB* gene of *L. biflexa* was expressed and a SOD activity could be detected into *L. interrogans* (Fig. S3, Fig. 1D). Surprisingly, the presence of an active SOD impaired *L. interrogans* growth in the presence of superoxide (Fig. 1E). We ratiocinated that the introduction of an active SOD in *L. interrogans* would increase the cytoplasmic H_2_O_2_ content produced consequently of superoxide dismutation. *L. interrogans* only have one catalase (KatE) located in the periplasm^6,12^ (Fig. 1B). We therefore tested whether the presence of a cytoplasmic catalase (KatG), together with SOD, would improve the *L. interrogans* growth in the presence of superoxide. Expression of the *L. biflexa* catalase-encoding *katG* in the *sodB*-expressing *L. interrogans* strain rescued its ability to grow in the presence of superoxide (Fig. 1E). *KatG* expression reduced the lag phase and increased the growth rate of the WT strain, that does not express *sodB*, in the presence of superoxide. This difference in tolerating superoxide could not be explained by a general fitness difference as all strains exhibited comparable growth in the absence of superoxide. Of note, expressing *L. biflexa sodB* did not significantly affect *L. interrogans* virulence (Fig. S3). These findings demonstrate that the presence of a SOD in *L. interrogans* does not improve the fitness in the presence of superoxide unless a cytoplasmic H_2_O_2_ detoxification system is provided. This supports a scenario whereby peroxidases and/or catalases have evolved in pathogenic *Leptospira* spp. accordingly with the loss of a cytoplasmic SOD.

### Long-lasting adaptation of *L. interrogans* to superoxide stress

We tested whether pathogenic *Leptospira* spp., despite the absence of a SOSE, have evolved alternative defense mechanisms and could adapt to superoxide. As previously reported^13–15^, when WT *L. interrogans* are cultivated in the presence of paraquat, bacterial growth resumes after a ∼10 day-lag phase (Fig. S4, cycle 1). Exponentially growing *L. interrogans* in the presence of paraquat were used to inoculate fresh EMJH with and without paraquat (Fig. S4, cycle 2). This superoxide-selected population exhibited a three-fold reduced lag phase (∼3.5 days) in the presence of paraquat compared to that not first cultivated in the presence of paraquat (Fig. S4, compare cycles 1 and 2).

To characterize the bacterial population selected under this condition, adaptation was repeated using a clonal population. A single clone was used to inoculate EMJH medium and two consecutive exposures to paraquat were performed as described above (Fig. 2A, cycles 1 and 2). The bacterial population selected in the presence of paraquat at cycle 1 exhibited, as before, a reduced lag phase when cultivated in the presence of superoxide (Fig. 2B, cycle 2). When this culture was propagated for 5 consecutive passages in the absence of superoxide, the bacterial population still exhibited a reduced lag phase (∼2.5 days) when cultivated in the presence of superoxide (Fig. 2B, cycle 7). We further propagated the superoxide-adapted population and the original culture for 39 consecutive passages (cycle 46, ≈94 generations) in the absence of paraquat. We observed that the superoxide-adapted population still conserved its adaptive phenotype in the presence of paraquat (Fig. S4, cycle 46). The superoxide-adapted population did not exhibit any fitness defect when compared to the non-adapted original population grown in EMJH in the absence of superoxide, nor it did exhibit a higher fitness in the presence of H_2_O_2_ (Fig. 2B, Fig. S4). Altogether, these findings demonstrate that *L. interrogans* can acquire a specific long-lasting adaptation to superoxide at no fitness cost in the absence of a SOSE.

**Figure 2.**
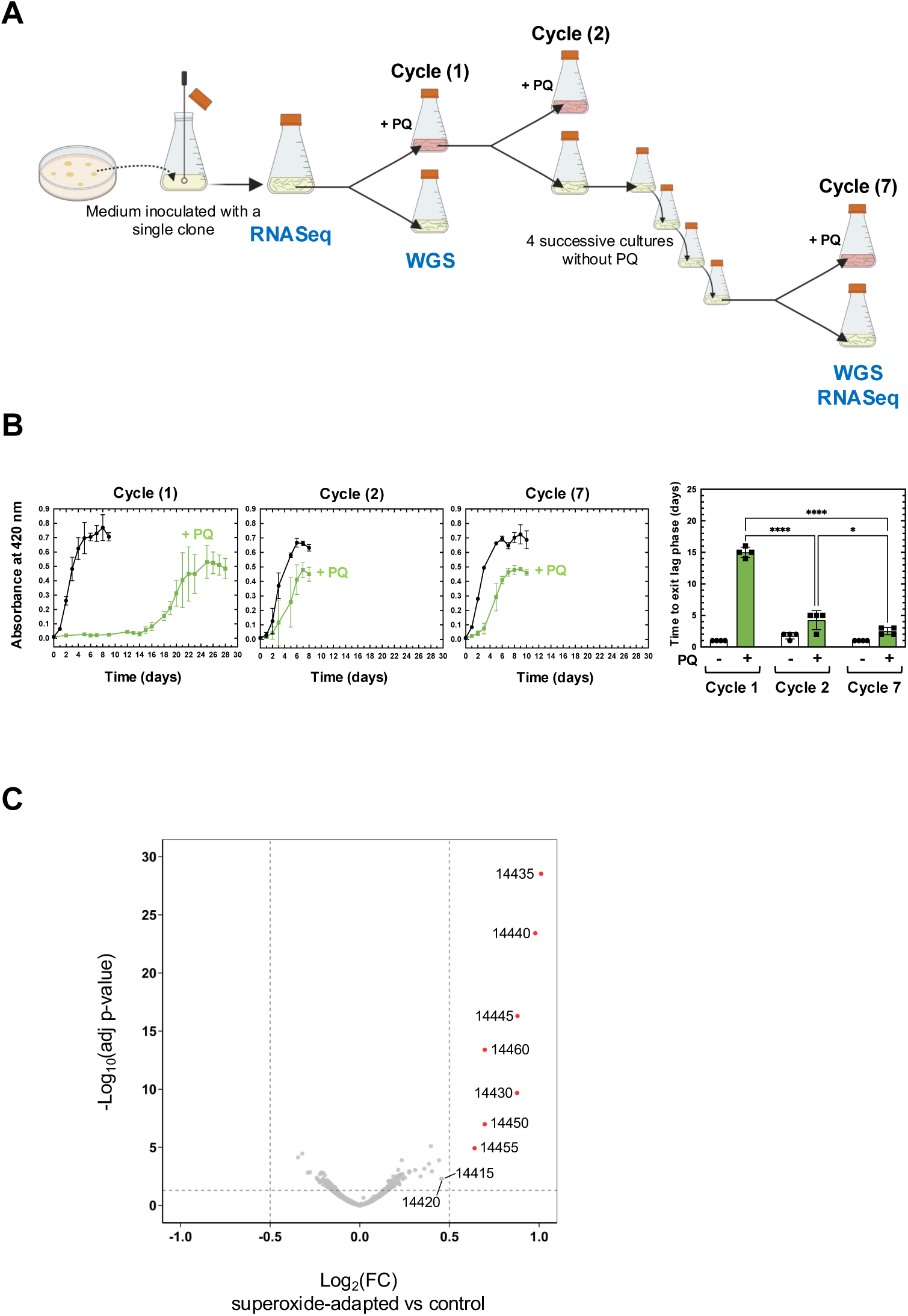
Adaptation of pathogenic *Leptospira* spp. to superoxide. (A) Schematic representation of the pipeline for experimental adaptation of *L. interrogans* to superoxide. The adaptation was performed by cultivating *L. interrogans* in the presence of 2 μM paraquat (PQ). The cycle numbers are the number of *in vitro* passages. The steps where whole genome sequencing (WGS) and RNA-seq had been performed are indicated. (B) Growth curve of WT at different stages of the adaptation in the absence (black circle) or presence (green square) of 2 μM paraquat (PQ). Growth was assessed by measure of absorbance at 420 nm. Time to exit lag phase (as defined by the first day where the cultures exhibit an OD≥0.05) was plotted for each cycle, with white bars (circle) and green bars (square) corresponding to growth in the absence or presence of paraquat, respectively. Data are mean and SD of four independent experiments. ****, p-value<0.0001; *, p-value=0.012. (C) Volcano representation of the DEGs in the superoxide-adapted strain. Differential expressions are expressed as Log_2_FC of the adapted strain (at cycle 7) versus the control strain. Significantly upregulated ORFs (cutoffs of Log_2_FC>0.5, adj. p-value<0.05 indicated by vertical and horizontal dashed lines) are represented in red and labelled with ORF number according to UP-MMC-NIID LP strain^41^. Please note that there is no significantly downregulated ORF.

To identify the mechanisms underlying this adaptation, the genome of the superoxide-adapted population at cycle 7 (Fig. 2A) was sequenced and compared with that of the non-adapted population at cycle 1. Short- and long-read whole genome sequencing demonstrated the absence of any single nucleotide polymorphisms (SNPs), indel, or genome rearrangements that could explain the adaptation (Table S3).

Comparative RNA-seq analysis indicated that only 11 genes were upregulated in the superoxide-adapted strain (0.4<Log_2_FC<1, *padj*<0.02) (Table S4). These genes encode poorly characterized proteins of the same locus (LIMLP_14415-14460), including an AsrR transcriptional regulator, a DoxX-like protein (LIMLP_14420), three START domain-containing proteins (LIMLP_14430-14440), a DHFR domain-containing protein (LIMLP_14455), and an MFS transporter (LIMLP_14460) (Fig. 2C, Fig. S5). Thus, emergence of adaptation to superoxide in *L. interrogans* is independent of permanent genome modifications and may be due to inheritable differential gene expression.

### The transcriptomic and proteomic responses to superoxide suggest a metabolism reprograming

To identify superoxide-induced mechanisms in pathogenic *Leptospira*, RNA-seq and mass spectrometry analyses were performed on *L. interrogans* exposed to 10 and 50 μM of paraquat. These conditions were chosen as they do not lead to more than 22% of bacterial death (Fig. S6). 18 and 139 ORFs were differentially expressed in the presence of 10 and 50 μM of paraquat, respectively (|Log_2_FC|>1, *padj*<0.05) (Fig. 3A, Tables S5-S7), showing a dose-dependent effect.

**Figure 3.**
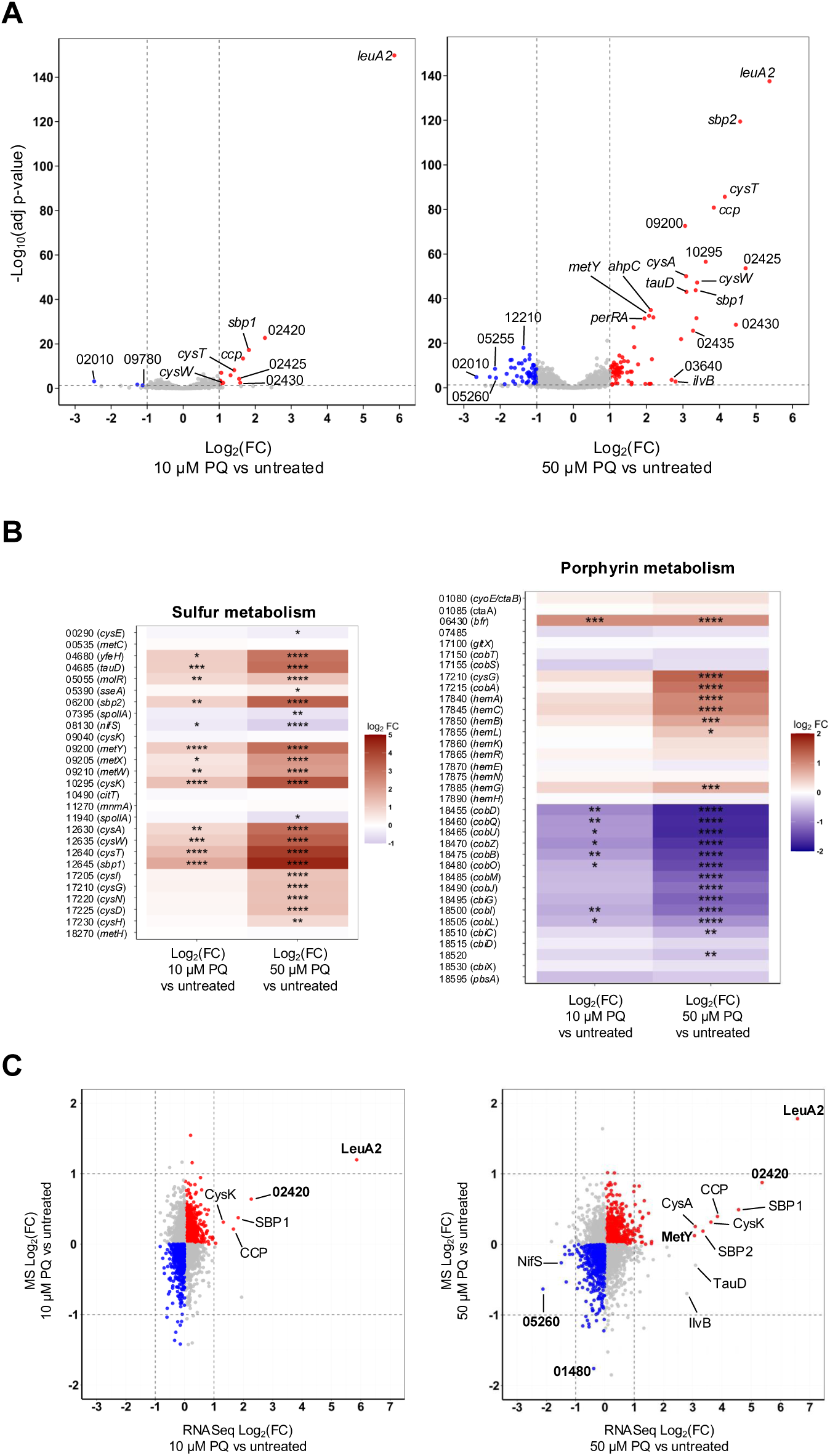
Transcriptomic and proteomic response of *L. interrogans* to superoxide. (A) Volcano representation of the DEGs in the presence of 10 (left panel) or 50 μM (right panel) paraquat (PQ). Differential expressions are expressed as Log_2_FC of PQ-treated versus untreated strain. Significantly downregulated and upregulated ORFs (cutoffs of |Log_2_FC|>1, adj. p-value<0.05 indicated by vertical and horizontal dashed lines) are represented in blue and red, respectively. Selected genes are labelled by gene name or ORF number according to UP-MMC-NIID LP strain^41^. (B) Heat map of DEGs related to sulfur (left panel) or porphyrin (right panel) metabolism in the presence of 10 or 50 μM PQ. Gene names and number (according to UP-MMC-NIID LP strain) are indicated on the left. The heat map color from blue to red indicates low to high Log_2_FC. *: adj. p-value<0.05, **: adj. p-value<0.01, ***: adj. p-value<0.001, ****: adj. p-value<0.0001 (C) Correlation between DEGs (obtained by RNASeq) and differentially produced proteins (obtained by mass spectrometry (MS)) in the presence of 10 (left panel) or 50 μM (right panel) PQ. Blue and red dots indicate down- and upregulated ORFs, respectively. Factors that are significantly differentially expressed in both techniques are highlighted in bold.

Several genes encoding oxidative stress-related factors were upregulated in the presence of paraquat. Notably, genes encoding the catalase KatE, the peroxiredoxin AhpC1, the cytochrome C peroxidase CcpA, as well as the two peroxide stress regulators PerRA and PerRB, exhibited an increased expression upon exposure to paraquat (Table S6). This indicates that paraquat triggers an oxidative stress in *Leptospira*.

Importantly, differentially expressed genes (DEGs) were enriched in metabolic pathways (Fig. S6). Remarkably, the most up-regulated gene at both concentrations encodes a 2-isopropylmalate synthase (*leuA2*), an enzyme that catalyzes the first step of leucine biosynthesis pathway (Fig. 3A, Fig. S7).

In addition, many genes related to import and metabolism of sulfur-containing molecules were significantly upregulated in the presence of paraquat (Fig. 3A-B, Fig. S7, Table S6). Genes encoding components of an ABC sulfate import complex (*sbp1, sbp2*, *cysAWT*) and enzymes of the assimilatory sulfate reduction that catalyze the conversion of sulfate (SO_4_^2-^) to sulfide (H_2_S) (i.e. *cysDNHI*), all exhibited increased expression in the presence of paraquat. Genes of the siroheme biosynthesis pathway (*cysG* and *cobA,* encoding a siroheme synthase and uroporphyrinogen-III C-methyltransferase, respectively) were as well upregulated by paraquat. Siroheme is an essential cofactor of the sulfite reductase CysI, necessary to reduce sulfite into sulfide. Another source of sulfite in bacteria is the sulfur-containing amino acid taurine.

LIMLP_04680, that encodes a putative bile acid:sodium symporter for taurine uptake (homolog to *Escherichia coli yfeH*) and the adjacent gene *tauD*, that encodes a putative taurine dioxygenase catalyzing the release of sulfite from taurine, were upregulated upon exposure to paraquat (Fig. 3B, Fig. S7, Table S6).

The assimilatory sulfate reduction pathway allows the incorporation of sulfide into cysteine and methionine. The reaction of H_2_S and O-acetyl L-homoserine leads to synthesis of L-cysteine and is catalyzed by the cysteine synthase CysK. The reaction of H_2_S with O-acetyl L-serine, that produces L-homocysteine, is catalyzed by a homocysteine synthase MetY. Both *cysK* and *metY* exhibited increased expression in the presence of paraquat (Fig. 3B, Fig. S7, Table S6). However, the gene encoding the methionine synthase MetH that converts homocysteine into methionine is not differentially expressed in the presence of paraquat (Table S5).

Nearly 50% of down-regulated genes encoded proteins with unknown function (Table S7). Others encoded factors of different metabolic pathways, including ATP (*atpGDC*) and cobalamin (*cobDQUZBOMJIL*, *cbiGCD*) biosynthesis (Fig. 3B, Table S7).

Analysis of the proteome in the same conditions confirmed the significant upregulation of LeuA2, LIMLP_02420-encoded protein and the downregulation of MauG (encoded by LIMLP_05260) (Fig. 3C, Table S8). In addition, several factors of the sulfur assimilatory pathway (Sbp1, Sbp2, CysA, CysN, CysD, CysH, CysK, MetY) had an increased cellular content, although with a Log_2_FC lower than 1 (Fig. 3C, Table S8).

Altogether, these findings indicate that, in the absence of SOSEs, the paraquat-induced oxidative stress triggers a metabolic reprogramming toward leucine and sulfur-containing amino acids (Cys and Met) biosynthesis pathways.

### The superoxide and peroxide-induced transcriptional response are partially distinct

Superoxide exposure triggers the upregulation of several genes of the PerRA regulon and H_2_O_2_ regulon, i. e. *katE*, *ahpC1*, *ccpA*, probably because its reduction leads to H_2_O_2_ production. To determine to what extent the superoxide and peroxide transcriptional responses overlap, we compared DEGs in the presence of paraquat and H_2_O_2_. We previously determined that 505 ORFs of *L. interrogans* are significantly differentially expressed in the presence of 1 mM H_2_O_2_^14^ (Table S9). 84% (*N=*422) of the H_2_O_2_ stimulon is not shared with the superoxide stimulon, and 40% (*N*=83) of the 137 DEGs upon exposure to 50 μM paraquat are not deregulated by H_2_O_2_. *KatE*, *ahpC1*, *ccpA*, *perRA*, *perRB*, *clpB*, the heme and cobalamin biosynthesis genes were deregulated with both ROS (Table S9). Therefore, a subset of the upregulation observed upon exposure of superoxide is probably due to the presence of H_2_O_2_ produced from the reduction of superoxide. However, the upregulation of genes of the sulfate assimilatory pathway and *leuA2* is greater upon paraquat exposure, suggesting that their upregulation is mostly due to specific superoxide-mediated stresses.

### Distinct differential expression of the *leuA* paralogs in the presence of superoxide

*LeuA2* (LIMLP_15720) is the most upregulated gene upon exposure to paraquat. The *L. interrogans* genome possesses two *leuA* paralogs, i.e. *leuA1* (LIMLP_08570) and *leuA2* (LIMLP_15720). Both ORFs exhibit the canonical catalytic domains of a 2-isopropylmalate synthase, however, LeuA2 lacks the C-terminal domain responsible for an allosteric inhibition by leucine (Fig. 4A, Fig. S8)^16^. Unlike LeuA2, LeuA1 is not differentially expressed in the presence of paraquat (Fig. 4B). Genus-wide comparative analysis revealed that LeuA1 and LeuA2 homologs contain 6 and 2 cysteine residues on average, respectively (Fig. 4C). Moreover, LeuA1 was predicted to be 3 times more redox-sensitive than LeuA2 (Fig. 4C). Despite leucine being the most frequent amino acid in *L. interrogans*, its abundance was not affected by the paraquat treatment (Fig. S8, Fig. S9). Thus, LeuA2 is the only factor of the leucine biosynthesis pathway that is upregulated by superoxide.

**Figure 4.**
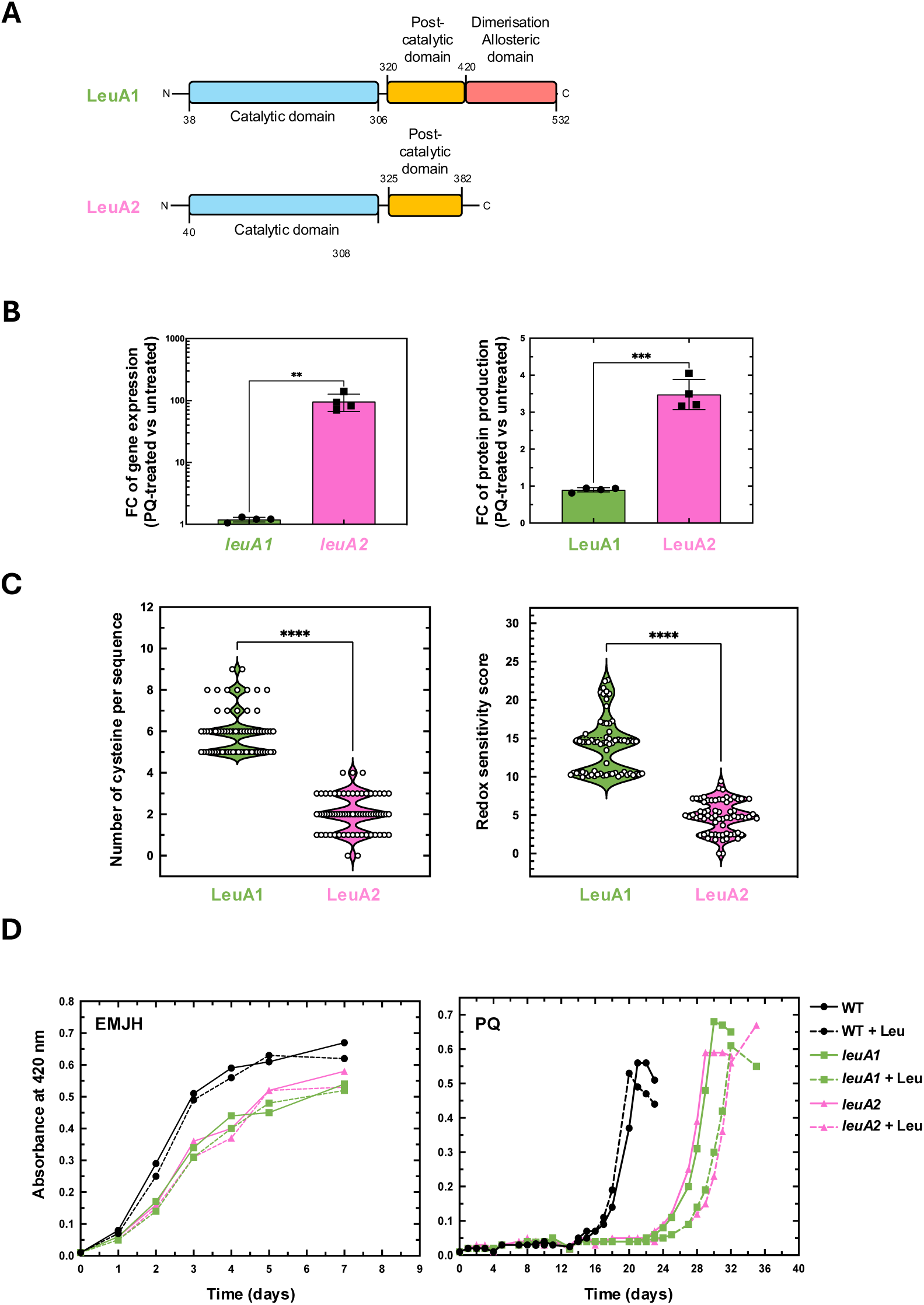
Role of LeuA1 and LeuA2 in the *L. interrogans* fitness in the presence of superoxide. (A) Schematic representation of the domain organization of LeuA1 (LIMLP_08570) and LeuA2 (LIMLP_15720) 2-isopropylmalate synthases with the catalytic pyruvate carboxyltransferase HMGL-like domain in blue, post-HMGL-like domain in yellow and dimerization allosteric domain in red. (B) Gene (left panel) and protein (right panel) expression of LeuA1 (green bar, circle) and LeuA2 (pink bar, square) upon exposure to superoxide. Differential expressions are expressed as FC (50 μM PQ-treated versus untreated). Data are mean and SD of four independent experiments. **: p-value<0.05, ***: p-value=0.0009. (C) Number of cysteine residues (right panel) and predicted redox-sensitive (as analyzed by IUPRED2A-redox^53^) score of LeuA1 (green) and LeuA2 (pink) in 68 *Leptospira* species. Each symbol represents a species. ****: p-value<0.0001. (D) Growth of *L. interrogans* WT (black circle), *leuA1* (green square) and *leuA2* (pink triangle) mutant strains in EMJH medium (left panel) or in the presence of 3 μM paraquat (PQ, right panel) with (dashed lines) or without (solid line) 2 mM leucine. Growth was assessed by measure of absorbance at 420 nm. Data are one replicate representative of 3 three independent biological replicates.

To determine whether LeuA1 and LeuA2 could have a role in tolerance to superoxide, the growth of WT, *leuA1* and *leuA2* mutant strains were assessed in the presence of paraquat. Inactivating *leuA1* or *leuA2* slightly impaired the growth of *L. interrogans* in the EMJH medium and severely compromised growth in the presence of paraquat (Fig. 4D). Addition of leucine did not improve the fitness of *L. interrogans* WT and mutant strains in the presence or absence of paraquat (Fig. 4D).

### Superoxide exposure leads to increase in cysteine oxidation

To assess the amino acid oxidation triggered by superoxide, proteinogenic amino acids detected in the proteome were quantified in *L. interrogans* upon exposure to paraquat. We determined that cysteine is the least abundant AA in the *L. interrogans* proteome (both *in silico* and by mass spectrometry analysis), and exposure to paraquat did not affect its abundance (Fig. S9, Fig. S10). However, exposure to 50 μM paraquat led to a significant increase in cysteine bi-oxidation (sulfinic acid) (Fig. 5A, Fig. S10).

**Figure 5.**
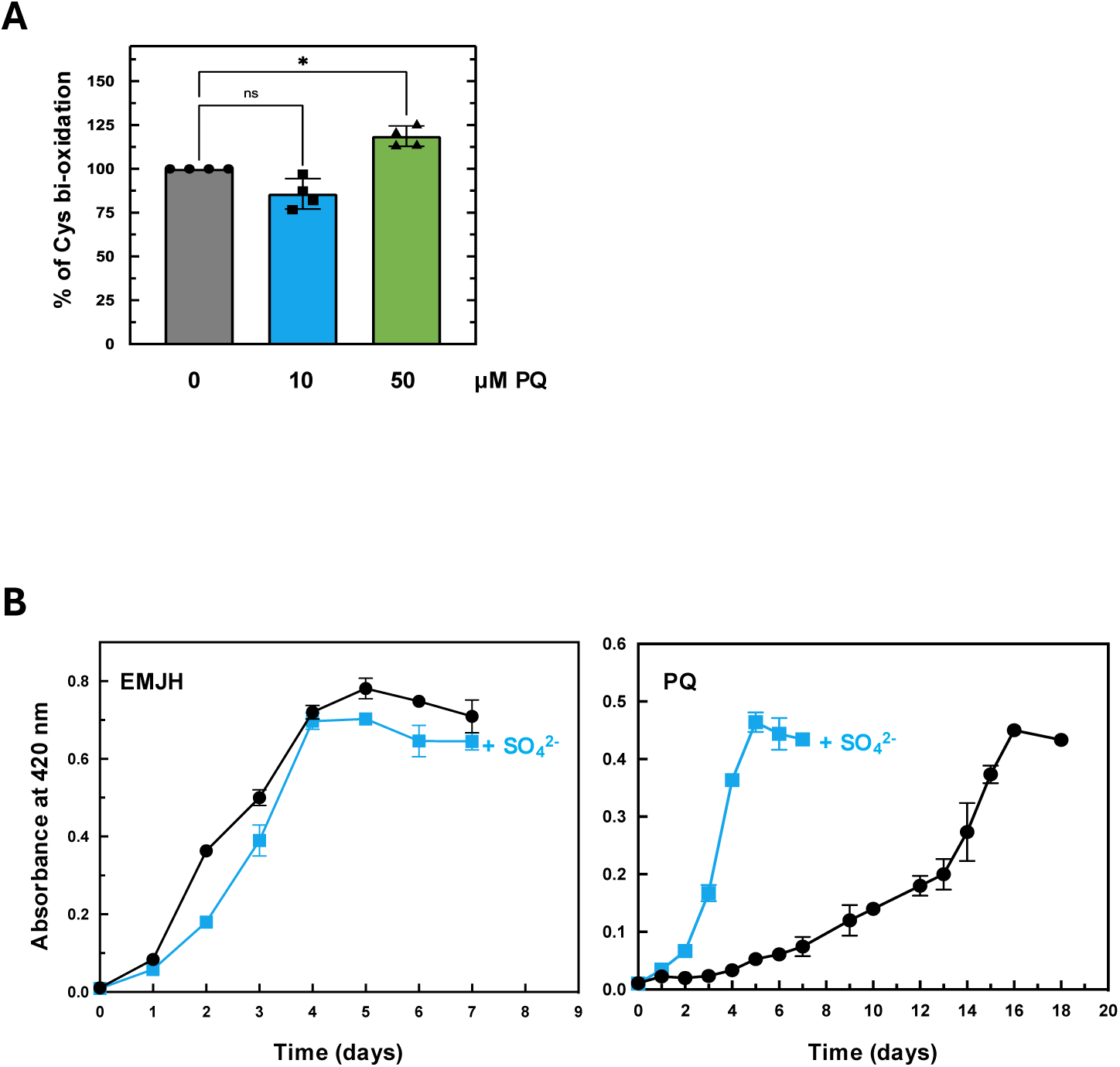
Addition of sulfate increases *L. interrogans* fitness in the presence of superoxide. (A) Mass spectrometry-based quantification of cysteine bi-oxidation (sulfinic acid) upon exposure to 0 (grey bar, circle), 10 (blue bar, square) or 50 (green bar, triangle) μM paraquat (PQ). Data are mean and SD of four independent experiments. *, p-value=0.0123 (B) Growth of *L. interrogans* WT strain in EMJH medium in the absence (left panel) or presence of 2 μM paraquat (PQ) (right panel) with (blue line) or without (black line) 20 mM Na_2_SO_4_. Growth was assessed by measure of absorbance at 420 nm. Data are mean and SD of three independent experiments.

Determining the proteome-wide oxidation ratios demonstrated that only a subset of proteins was oxidized in cysteine. DnaG primase (encoded by LIMLP_08540) was the protein exhibiting the highest significant increase in its cysteine sulfinic acid content (at Cys321) upon exposure to paraquat (Fig. S11, Table S8). Interestingly, DnaG oxidation correlated with a replication arrest when *L. interrogans* was exposed to superoxide (Fig. S11). Overall, downregulated genes and proteins were enriched in cysteine-containing proteins (Fig. S11) and almost none of the predicted Fe-S cluster-containing proteins exhibited increased expression upon exposure to paraquat (Fig. S11). In addition, exposure to paraquat did not significantly affect aconitase activity, used here as a proxy for assessing damage to iron-sulfur cluster-containing proteins (Fig. S11). Overall, this demonstrates that superoxide exposure causes cysteine oxidation without leading to Fe-S cluster inactivation.

### Sulfate increases adaptation to superoxide

The upregulation of several factors of the sulfate assimilatory pathway upon exposure to superoxide strongly suggests a role of this pathway for the adaptation to superoxide. To confirm this, metabolites (sulfate, sulfite, sulfide) and final product (cysteine) of the pathway were added to the culture medium. While the growth of *L. interrogans* in the presence of superoxide was not improved when sulfite (SO_3_^2-^), sulfide donors (Na_2_S, NaSH), cysteine or cystine (cysteine dimer) were added to the medium, the addition of sulfate did increase the ability of *L. interrogans* to grow in the presence of superoxide (Fig. 5B, Fig. S12). Exposure of *L. interrogans* to superoxide did not trigger an increase in sulfide (H_2_S) production (Fig. S12), indicating that H_2_S is not the final product of the sulfur assimilatory pathway upon paraquat exposure. Thus, we confirmed that sulfate assimilation improves adaptation of *L. interrogans* to superoxide.

## Discussion

In the present study, we demonstrate that all P clade *Leptospira* species lack SOD. No superoxide removal activity could be observed upon exposure to superoxide (Fig. S3), confirming that the life-threatening pathogen *L. interrogans* is devoid of an alternative superoxide detoxification machinery. Importantly, we reveal that the SOD present in the LLCA was maintained in saprophytes but specifically lost by pathogens. Our findings indicate that SOD loss could have occurred stepwise, as within the S2 clade, leading to P species devoid of any *sod*. In addition, maintaining a SOD activity in *Leptospira* species requires the concomitant presence of cytoplasmic H_2_O_2_ detoxification enzymes, such as KatG. What could have driven SOD loss? Since the main function of this enzyme is to rapidly eliminate the toxic superoxide produced during aerobic metabolism, it is tempting to propose that the aerobic metabolism of P-clade *Leptospira* species is reduced in the low oxygenated environment encountered within host tissues. Alternatively, SOD loss could have been driven by a reduced host-produced superoxide accumulation at the site of colonization during infection.

The absence of a SOSE in bacteria is rare (estimated at 9-13%), as demonstrated here and in a recent study^17^. The loss of a SOD is unforeseen in pathogenic bacteria as this enzyme has proven to be necessary for virulence of a variety of bacteria^18–20^ and some pathogens have even acquired additional extracellular SOD to fight host-derived superoxide^21,22^. Furthermore, our finding that SOSE-deficient organisms are equally found among aerobes and anaerobes suggests that aerobicity does not determine the prevalence of SOSEs. Overall, this challenges the universality of the oxygen toxicity theory and the essentiality of superoxide scavenging activity for aerobes.

Despite the absence of any SOSE, we successfully selected a long-lasting superoxide-adapted population after a unique exposure to superoxide. This adaptation, inherited for at least 94 generations, was independent of any mutation and at no fitness cost. A differentially expressed gene cluster could be identified in the superoxide-adapted strain and these uncharacterized genes may be involved in the adaptation. However, the modest upregulation of this cluster cannot solely explain the long-lasting adaptation to superoxide. Inheritable epigenetic modifications could also participate in the adaptation to superoxide. Interestingly, this adaptation was characterized by a reduced lag phase rather than a decrease in the doubling time, suggesting that the superoxide-adapted population is more efficiently prepared for the transcriptional and metabolic reprogramming necessary to exit the lag phase. In a previous study, we had identified a PerR regulator in *L. interrogans* (PerRB) whose inactivation led to an increased fitness in the presence of superoxide^15^, corroborating that pathogenic *Leptospira* species possess alternative transcriptionally controlled defense mechanisms to compensate for the loss of SOD.

In this study, we demonstrate that the sulfate assimilatory reduction pathway is upregulated upon exposure to superoxide. The importance of this pathway is experimentally confirmed by showing that sulfate improves the fitness of *L. interrogans* in the presence of superoxide. This pathway leads to the production of H_2_S, a molecule with antioxidant properties^23^, and to sulfur-containing amino acids (Cys and Met). We could not detect H_2_S production in *L. interrogans*, even in the presence of paraquat (Fig. S12). It is therefore unlikely that the upregulation of the sulfate assimilatory reduction pathway leads to the accumulation and protective effect of H_2_S. In addition, the facts that a *metY* mutant did not exhibit any fitness defect with superoxide (Fig. S12) and that *metH* is not deregulated with superoxide make it very unlikely that methionine participates in defense against superoxide. The upregulation of the cysteine synthase CysK strongly suggests that the purpose of the superoxide-triggered sulfate assimilatory reduction pathway is cysteine synthesis. Cysteine could either be used for the synthesis of the redox buffer glutathione or act as thiol protectant through S-thiolation, as demonstrated in *Staphylococcus aureus* and *Bacillus subtilis*^24,25^ (Figure 6A-B). Cysteines could also be incorporated in newly synthesized polypeptides to replace damaged proteins containing irreversibly oxidized cysteines (Figure 6A-B). Lastly, cysteines catabolism could contribute to iron-sulfur cluster assembly (Figure 6A-B). However, no genes involved in iron-sulfur cluster assembly, glutathione biosynthesis, or cysteine catabolism were upregulated in our conditions. While the fate of cysteine upon exposure to superoxide is yet to be determined, we demonstrate that the sulfate assimilatory reduction pathway plays a central role in adaptation to superoxide. Consistent with this, CysK is among the 4 most abundant proteins in *L. interrogans*^26^, thereby suggesting that pathogenic *Leptospira* compensated the absence of SOSEs by maintaining high levels of cysteine production.

**Figure 6.**
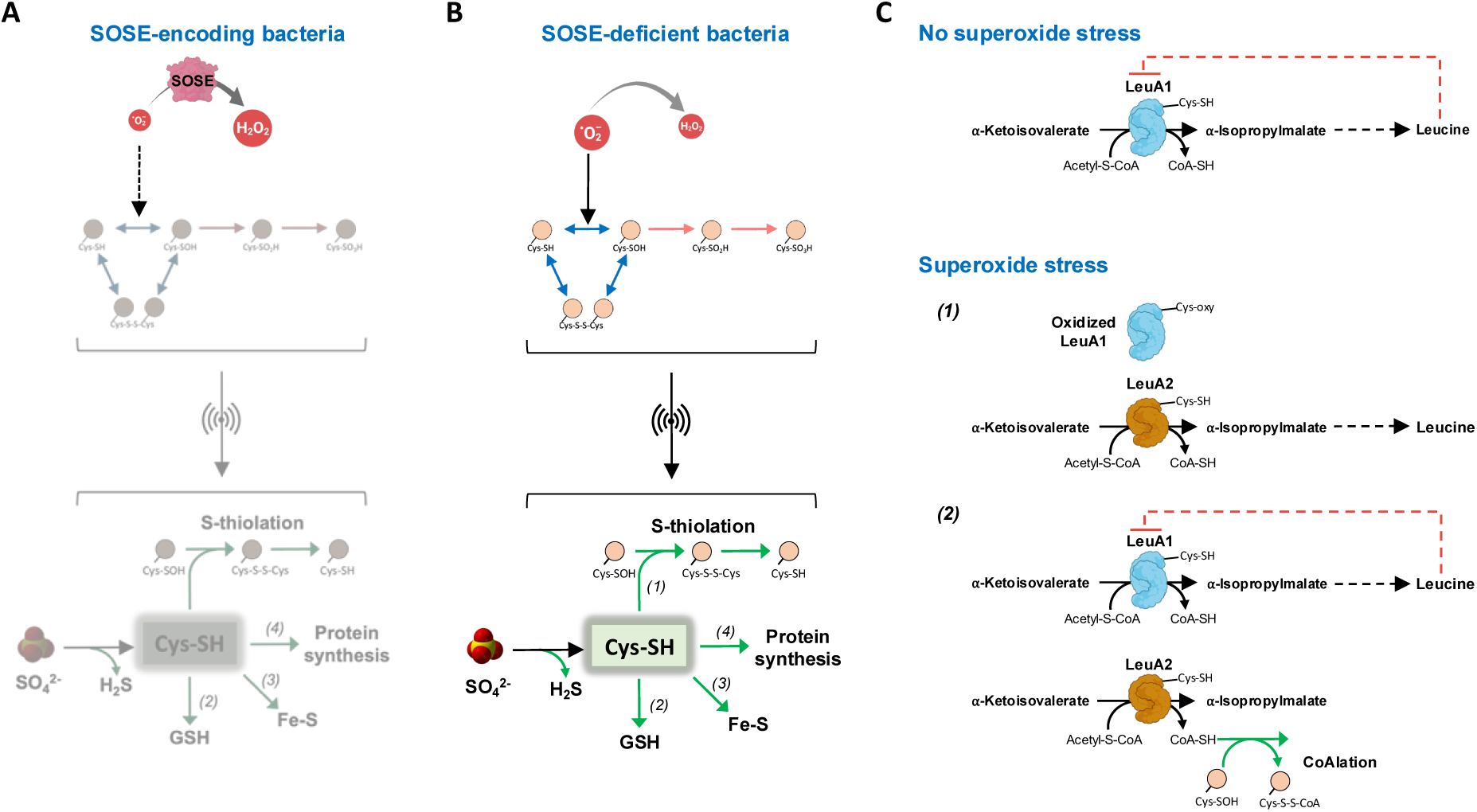
Model for the superoxide stress response in SOD-deficient bacteria. (A) In SOSE-containing organisms, rapid reduction of superoxide into hydrogen peroxide prevents cysteine oxidation and upregulation of the sulfur assimilatory pathway. (B) In SOSE-deficient organisms, such as *L. interrogans*, the low rate of superoxide reduction favors reversible (disulfide bond and sulfenic acid) and irreversible (sulfinic and sulfonic acids) cysteine oxidation, leading to protein denaturation and damage. The upregulation of the sulfur assimilatory pathway might result in an increased cysteine cellular content. Cysteine could contribute to resistance to oxidative damage by protecting thiol groups of cysteine residues from oxidation through thiolation (1), by being used for glutathione (GSH) synthesis (2), by providing sulfur for iron-sulfur clusters (3), by being incorporated in newly synthesized proteins to replace irreversibly damaged protein during oxidation (4). H_2_S, an intermediate metabolite of the sulfur assimilatory pathway, might also have antioxidant properties. Accumulation of oxidized cysteines might signal the upregulation of the sulfur assimilatory pathway. (C) Model for the role of LeuA2 in the superoxide stress response. In normal growth condition, LeuA1 is the isopropylmalate synthase that catalyzes the first step of leucine synthesis pathway (represented on the top). The inhibition of LeuA1 by leucine is represented by a dashed red arrow. Two mechanisms can explain LeuA2 upregulation upon superoxide stress. In a first mechanism, LeuA1 is oxidized and LeuA2 production compensates for the inactivation of LeuA1 (1). In another mechanism, LeuA2 is produced during oxidative stress to trigger accumulation of CoASH that protects oxidized cysteine by CoAlation (2). In the latter case, LeuA1 could be among the CoAlated proteins. LeuA2 lacks the C-terminal regulatory domain and is not inhibited by leucine.

The most upregulated factors upon exposure to superoxide was the 2-isopropylmalate synthase that catalyzes the first step of leucine biosynthesis encoded by *leuA2*. Of the two *leuA* paralogs present in *Leptospira* spp., only *leuA2* was upregulated by superoxide. In the absence of oxidants, LeuA1 is 3.7 times more abundant than LeuA2^26^, and LeuA2 does not contain the C-terminal domain responsible of allosteric inhibition by leucine^16^. It is very likely that LeuA1 is the enzyme in charge of maintaining leucine homeostasis in the absence of oxidants, while the specific upregulation of LeuA2 by superoxide would allow to compensate for the oxidative damage to LeuA1 (Figure 6C, mechanism 1). This hypothesis is supported by the fact that LeuA1 homologs, which contain more cysteine than LeuA2s, are predicted to be more redox sensitive. Another hypothesis is that the conversion of 2-keto-isovalerate and acetyl-CoA to 2-isopropylmalate, when catalyzed by LeuA2, would lead to coenzyme A (CoA-SH) accumulation, allowing protection of cysteine thiols from oxidation by CoAlation^27,28^ (Figure 6C, mechanism 2).

A classical interpretation of the role of leucine and cysteine in superoxide stress stems from the conditional auxotrophy of *E. coli sod* mutants for several amino acids under aerobic conditions^29,30^. This was explained by inactivation of Fe-S cluster of enzymes involved in amino acids biosynthesis^31,32^. In our conditions, addition of cysteine or leucine did not improve the fitness of *L. interrogans* in the presence of superoxide, and aconitase activity was not altered by exposure to superoxide. While we cannot exclude that higher paraquat doses would lead to Fe-S cluster inactivation in *L. interrogans*, we speculate that SOSE-deficient organisms, such as pathogenic *Leptospira* spp., have evolved higher Fe-S cluster resistance from oxidative damage. To conclude, by exploring adaptation of a naturally SOSE-deficient species, like pathogenic *Leptospira*, to superoxide, we provide evidence challenging the classical dogma of the oxygen toxicity. We demonstrate that these aerobes, which have lost their SOD, are still capable of adaptation and have evolved a metabolic trade-off to compensate for the absence of an enzymatic superoxide detoxification activity.

## Methods

### Bacterial strains and growth condition

*L. interrogans* serovar Manilae strain L495 and transposon mutant strains (see Table S10 for a complete description of the transposon mutants used in this study) were grown aerobically at 30°C in Ellinghausen-McCullough-Johnson-Harris medium (EMJH)^33^ with shaking at 100 rpm or in EMJH solid agar medium for 1 month until appearance of colonies. ΙΙ1 and β2163 and *E. coli* strains were cultivated at 37°C in Luria-Bertani medium with shaking at 37°C in the presence of 0.3 mM thymidine or diaminopimelic acid (Sigma-Aldrich), respectively. Spectinomycin was added to the media at 50 µg/ml when needed. When indicated, the superoxide-generating compound paraquat (1,1′-Dimethyl-4,4′-bipyridinium-2,2′,3,3′,5,5′,6,6′-d8 dichloride), H_2_O_2_ (Sigma-Aldrich), sodium sulfate (Na_2_SO_4_), sodium bisulfite (NaHSO_3_), sulfide donors (Na_2_S, NaSH), leucine, cysteine, cystine (diluted in Tris-HCl), or Tris-HCl were added to the medium. Bacterial growth was followed by measuring the absorbance at 420 nm. The number of generations was estimated under the assumption that the average generation time of *L. interrogans* is 20 hours and that each passage corresponds to 48 hours of exponential growth.

### Expression of L. biflexa sodB and katG in L. interrogans

Heterologous expression of *sodB* in *L. interrogans* was performed by PCR amplification of LEPBIa0027 (*sodB*) from genomic DNA of *L. biflexa* serovar Patoc strain Patoc 1 using the primers sodB_F and sodB_R (Table S11). The PCR product was cloned into the pMaGRO vector at NdeI/XbaI restriction sites^34^. The resulting plasmid (pMAGRO::*sodB*) was named pSGH1. For heterologous expression of *katG*, a synthetic construct fusing the promoter of *lipL32* until its transcription start site to *katG* (LEPBIa2495) and flanked by BamHI restriction sites was obtained through synthesis (GeneArt, ThermoFisher). Cloning of the fusion was performed by restriction-ligation with BamHI in both empty pMaORI^35^ and pMaGRO::*sodB* (pSGH1). The resulting pMaORI::*katG* and pMaGRO::*sodB*-*katG* plasmids were named pSGH11 and pSGH10, respectively. All plasmids were verified by whole plasmid sequencing with Nanopore long-read technology before conjugation (Plasmidsaurus Inc, Arcadia, California) and were introduced in *L. interrogans* by conjugation, as previously described^36^. Conjugants were selected on solid EMJH agar containing spectinomycin (50 µg/ml) and correct plasmid acquisition was further verified by PCR amplification using the primers pMaORI-A/Mao2 and maoEco1/maoEco2 (Table S11). The WT *L. interrogans* containing empty pMaORI as control was obtained from previous studies^13^.

### RNA purification

Virulent *L. interrogans* serovar Manilae strain L495 with less than three *in vitro* passages were used in this study. Four independent biological replicates of exponentially grown WT *L. interrogans* strain were incubated in the presence or absence of 10 μM and 50 μM paraquat for 60 min at 30°C. Each sample was divided in two. One part was used for RNA purification and the other part was used for mass spectrometry analysis (see below). For the analysis of the superoxide-adapted strain, three independent biological replicates of control (non-adapted) or adapted strains were used. Harvested bacteria were resuspended in 1 ml TRIzol (ThermoFisher Scientific) and stored at −80°C. Nucleic Acids were extracted with chloroform and precipitated with isopropanol as described elsewhere^37^. Contaminating genomic DNA was removed by DNAse treatment using the RNAse-free Turbo DNA-free turbo kit (ThermoFisher Scientific) as described by the manufacturer. The integrity of RNAs (RIN>7) was verified by the Agilent Bioanalyzer RNA NanoChips (Agilent technologies, Wilmington, DE).

### RNA Sequencing

Libraries were built using a Illumina Stranded Total RNA Prep with Ribo-Zero library kit (Illumina, USA) following the manufacturer’s protocol. Quality control was performed on an Agilent BioAnalyzer. Sequencing was performed on Illumina’s platforms NextSeq500 to obtain 70 base single-end reads. The RNA-seq analysis was performed with Sequana 0.16.3^38^. We used the RNA-seq pipeline 0.19.1 (https://github.com/sequana/sequana_rnaseq) built on top of Snakemake 7.32.4^39^.

Briefly, reads were trimmed from potential adapters using Fastp 0.23.2^40^ then mapped to the *L. interrogans* serovar Manilae strain UP-MMC-NIID LP assembly^41^ (GCF_001047635.1_ASM104763v1) using Bowtie 2.4.5^42^. FeatureCounts 2.0.1^43^ was used to produce the count matrix, assigning reads to features using the corresponding genome annotation with strand-specificity information. Quality control statistics were summarized using MultiQC 1.17. Statistical analysis on the count matrix was performed to identify differentially regulated genes. Clustering of transcriptomic profiles were assessed using a Principal Component Analysis (PCA). Differential expression testing was conducted using DESeq2 library 1.34.0^44^ indicating the significance (Benjamini-Hochberg adjusted p-values, false discovery rate FDR < 0.05) and the effect size (fold-change) for each comparison. Differential expressions were expressed as logarithm to base 2 of fold change (Log_2_FC).

### *In silico* identification of SOD and SOR in bacterial genome databases

The SOR protein sequence from *Pyrococcus furiosus* (UniProt Q8U1K9) and the SOD protein sequence from *L. biflexa* (UniProt Q1EMH7) were used as queries for a search with BLAST v2.13.0^45^ against two databases constructed using the 1110 established bacterial pathogens infecting humans database^9^. Significant hits with e-value cutoff of 0.01 were extracted and aligned with MAFFT v7.467 (L-INS-i algorithm)^46^ to generate an HMM profile subsequently used to scan the databases for missing hits with HMMer v3.3.1^47^ with an e-value cutoff of 0.01. Presence of a SOD or SOR was estimated positive for all proteomes containing hits with an e-value cutoff lower than 0.01 for any of the two searches (BLAST or HMMer). To rule out artefacts due to lack of representativeness of the genome chosen within the species, all SOD/SOR double-negative species were double-checked by running BLASTp with the same queries against all RefSeq curated genomes available within each TaxID in the NCBI BLASTp website^48^. Positive hits for this last search were manually corrected in the final table. Aerobicity for the final list of SOD/SOR double-negative species was determined using literature screening.

### Distribution of *sod*, *katE* and *katG* in *Leptospira* spp

The SodB protein (LEPBIa0027), the KatE protein (LIMLP_10145) and the KatG protein (LEPBIa2495) sequences were used as query for a BLAST v2.13.0 search^45^ against the reference database of 68 *Leptospira* species^49,50^. Hits with an e-value ≤ 0.01 were retained and subsequently aligned by MAFFT v7.467 under the L-INS-i algorithm^46^. This alignment was used to compute an HMM profile to search for missing hits in the same database with HMMer v3.3.1^47^. All significant hits (e-value ≤ 0.01) were retained and aligned with MAFFT v7.467 under the L-INS-i algorithm and tree inference was performed with IQ-TREE v2.0.6 under the best-fitted model of evolution^51^. Putative orthologs were extracted from these phylogenies and plotted against a core-genome tree of *Leptospirales* obtained as described previously^12^ using the *ggtree* package for R v4.3.2^52^.

### SOD activity

For determination of SOD activity, 100 ml of exponentially growing *L. interrogans* or the respective *sodB*- and *katG*-encoding strains were harvested and resuspended in 800 µl of PBS and sonicated (3 cycles of 30 seconds) to prepare total protein extracts. Total protein extracts were then assayed for SOD activity using the Superoxide Dismutase Colorimetric Activity Kit (EIASODC, ThermoFisher) following the manufacturer’s recommendations.

### Redox-sensitivity score

Direct orthologs of LeuA1 (LIMLP_08570) and LeuA2 (LIMLP_15720) were searched as abovementioned and submitted to IUPRED2A with redox state option locally^53^. For the 68 *Leptospira* species, the redox sensitivity score was obtained by calculating the difference between the score under the redox-plus and the redox-minus profiles for all residues. Similar analysis was performed onto the 2-isopropylmalate synthase CimA (encoded by LIMLP_07785, 516 AA) as a control to ensure that the redox sensitivity score is independent of the protein length.

### Proteomic analyses

#### Sample preparation

*Leptospira* cultures (prepared as described for RNA extraction) were harvested and extensively washed with PBS. Pellets were resuspended in a lysis buffer (50 mM Tris-HCl pH 8.0, 2 mM EDTA, 5 mM DTT) and lysis was performed by sonication. 10 μg of total extract were loaded on a 12% SDS-PAGE. After 1 cm of migration in the resolving gel, proteins were fixed with acetic acid and ethanol, and stained by Colloidal Blue. In-gel protein digestion and extraction of tryptic peptides were performed as previously described^54^ with minimal variations. Briefly, 1 cm bands were excised from the SDS-polyacrylamide gel, previous to cysteine reduction and alkylation by incubation with 10 mM dithiothreitol (DTT) and 55 mM iodoacetamide (IAA), respectively. In-gel protein digestion was performed overnight at 37°C with sequencing grade trypsin (Promega) in a protease:protein ratio of 1:50 (w/w). Tryptic peptides were extracted from the gel with 60% ACN/0.1% trifluoroacetic acid (TFA) by two incubations of 1h at 30°C. The digestions were dried under vacuum and peptides were desalted using ZipTips C18 microcolumns (Merck Millipore). Desalted peptides were vacuum dried and resuspended in 0.1% formic acid (FA).

#### LC-MS/MS analysis

An UltiMate 3000 nanoHPLC system (Thermo Fisher Scientific) coupled to a Q Exactive Plus mass spectrometer (Thermo Fisher Scientific, USA) was used to perform LC-MS/MS analysis. Tryptic peptides were loaded into a precolumn (Acclaim PepMapTM 100, C18, 75 µm X 2 cm, 3 µm particle size) and separated in an Easy-Spray analytical column (PepMapTM RSLC, C18, 75 µm X 50 cm, 2 µm particle size) at 40°C using two mobile phases: 0.1% FA in water (A) and 0.1% FA in acetonitrile (ACN) (B). The separation gradient was from 1% to 35% B over 150 min and from 35% to 99% B over 20 min, at a flow rate of 200 nL/min. For data acquisition, the mass spectrometer was set in a positive mode using a top-12 data-dependent method, with an ion spray voltage of 2.3 kV and a capillary temperature of 250°C. The full MS scans were acquired from 200 to 2000 m/z with a resolution of 70000 at 200 m/z, an AGC target value of 1E6 and a maximum ion injection time of 100 ms. The precursors were fragmented in an HCD cell and acquired with a resolution of 17500 at 200 m/z, an AGC target value of 1E4 and a maximum ion injection time of 50 ms. For fragmentation, normalized collision energy (NCE) was used in steps of 25, 30 and 35. Dynamic exclusion time was set to 5 s. Each peptide sample was injected twice as technical replicates.

#### Mass spectrometry data analysis

PatternLab for Proteomics V software (PatternLab)^55^ was used to perform peptide spectrum matching and label-free quantitative analysis. MS data was searched against a target reverse database generated with PatternLab, including the *Leptospira interrogans* serovar Manilae strain UP-MMC-NIID-LP proteome (downloaded from NCBI, PRJNA287300, 20/09/2021) and the most common protein contaminants. For peptide identification m/z precursor tolerance was set at 40 ppm. Methionine oxidation, cysteine dioxidation, cysteine trioxidation and cysteine carbamidomethylation were defined as variable modifications. A maximum of 2 missed cleavages and 2 variable modifications per peptide were allowed. Search results were filtered by the PatternLab Search Engine Processor (SEPro) algorithm with a maximum FDR value ≤ 1% at protein level and 10 ppm tolerance for precursor ions. PatternLab’s Venn diagram statistical module was performed according to a Patternlab’s bayesian model to determine peptides and proteins uniquely detected in each biological condition using a probability value less than 0.05^56,57^. PatternLab’s TFold module was used to relatively quantify peptides and proteins present in both biological conditions by a spectrum count-based label-free quantification method. Peptides or proteins present in at least 5 biological replicates from the total of 8 were considered for TFold analysis. This module uses the Benjamini-Hochberg’s theoretical estimator to deal with multiple T-tests and it maximizes the number of identifications satisfying a fold change cutoff that varies with the p-values (BH q<0.05); while restricting false differential proteins mainly due to low abundance^58^.

### Statistics and reproducibility

Statistical analyses were performed with GraphPad Prism (version 10.3.0). All experiments were performed in at least three independent biological replicates. Unless otherwise stated, data represent mean and standard deviation of at least three independent biological replicates. All RNA-seq analyses were conducted using the Sequana RNA-seq pipeline (0.19.1) with published singularity/apptainer containers^59^, all of which are available on Zenodo via the Damona project (https://damona.readthedocs.io).

## Acknowledgments

We thank Elodie Turc, Laurence Ma, Georges Haustant, and Rania Ouazahrou of the Biomics Platform, C2RT, Institut Pasteur, Paris, France, supported by France Génomique (ANR-10-INBS-09) and IBISA, for the RNA-seq experiments. We also would like to thank Camille Zaniolo for her technical help in cloning *sodB* gene. We are grateful to Alexandre Giraud-Gatineau and Killian Coullin for critically reading this manuscript.

This work was supported by the PTR2019-310 grant (NB) from the Institut Pasteur, by National Institutes of Health grant P01 AI 168148 (MP & NB) and by FOCEM-COF 03/11 (AL & RD). SGH was a recipient of the Pasteur-Paris University PhD program.

