## Supplementary information for "Revisiting oxygen toxicity: evolution and adaptation to superoxide in a SOD-deficient bacterial pathogen"

SUPPLEMENTARY METHODS

### Phylogenomic analyses

SOD orthologs were searched in a database of 164 representative genomes of the phylum *Spirochaetes* following the approach described in the Methods section and used to infer a phylogeny with IQ-TREE v2.0.6 under the best-fitted model of evolution (WAG+I+G4)[1]. SOR orthologs were searched as described in the Methods section and both SOD and SOR spirochete orthologs were mapped to a core-genome phylogeny of *Spirochaetes* obtained using a concatenated alignment of 138 soft-core markers (identified with OMA v2.5.0) with IQ-TREE v2.0.6 under the best-fitted model of evolution (LG+F+I+R10)[1]. The rooting of the *Spirochaetes* tree was placed according to the previous analysis in Gupta et al. 2013[2], aerobicity was determined through a literature screening, and the final figures were generated using iTOL[3] or *ggtree* for R v4.3.2[4]. Locus analysis was performed using GeneSpy[5], and alignments were visualized with JalView v2.11.3.3[6].

### Whole genome sequencing

Genomic DNA for Illumina and Nanopore sequencing was extracted in four biological replicates for the adapted populations and the original clonal stock using the Maxwell 16 cell DNA purification kit (Promega).

### Variant calling

Variant calling and indel analysis of the Illumina sequencing data was performed as previously described in Zavala-Alvarado et al. 2021[7] using the *L. interrogans* serovar Manilae strain UP-MMC-NIID LP genome[8] as reference (accession numbers CP011931, CP011932, CP011933). Briefly, sequence reads were processed by fqCleaner 23.12 and aligned using Burrows-Wheeler Alignment tool (BWA mem 0.7.5a[9]). SNP and indel calling was performed with GATK2 following the Broad Institute best practices[10], and snpEff for variant annotation[11]. Comparison between VCF files was made with bedtools intersect[12].

Nanopore reads were aligned to the reference genome *Leptospira interrogans* serovar Manilae assembly (GCF_001047635_1_ASM104763v1) with minimap2 2.18-r1015[13] integrated in sequana mapper 1.2.1 workflow[14]. The bam files obtained were then analyzed using the sequana variant_calling 1.0.2 (https://github.com/sequana/variant_calling). Then, duplicated reads were ignored and aligned reads with a mapping quality score <30 were removed from further analysis. The workflow is then using Freebayes v1.2.0[15] (default options except for ploidy set to 10) for variant detection and statistics followed by SNPeff 5.1d[11] (with build options '-noCheckCds -noCheckProtein') for variants annotation with filtered VCFs (parameters: freebayes_score>20, frequency>0.1, min_depth>10, forward_depth>3, reverse_depth>3, strand_ratio>0.2). Quality control statistics were summarized using MultiQC 1.17[16]. Putative copy number variations (CNVs) were investigated using sequana coverage 0.16.11[17] with default parameters. Genomic rearrangements were assessed using Syri 1.6.3[18] and plotsr 1.1.1[19].

**Analysis of DEGs**

Gene ontology enrichment analysis was performed for all significantly deregulated genes under 50 µM paraquat (*padj*<0.05) with the ClueGO app v2.5.10[20] for Cytoscape v3.10.2[21]. Parameters were setup as follows: minimum GO level = 7, maximum GO level = 15, minimum percentage of genes = 50%, kappa score threshold = 0.4, GO fusion = false, GO group = true, two-sided hypergeometric test, p-value < 0.05.

**Replication measurements**

For replication measurements, 300 µl of bacterial culture were used to extract DNA using the Maxwell 16 cell DNA purification kit (Promega) at the indicated timepoints and genomic replication was quantified using primers against the *oriC* by quantitative PCR (Table S11).

Quantitative PCR was conducted with the SsoFast EvaGreen Supermix (Biorad) as previously described[22].

**SOD activity in the presence of paraquat**

Exponentially growing *L. interrogans* WT in EMJH medium were exposed or not to 50 µM paraquat for 1 hour. 50 ml of both cultures and 50 ml of exponentially growing *E. coli* K12 C600 (included as positive control) were harvested, resuspended in 800 µl of PBS and sonicated (3 cycles of 30 seconds) to prepare total protein extracts. Total protein extracts were then assayed for SOD activity using the Superoxide Dismutase Colorimetric Activity Kit (EIASODC, ThermoFisher) following the manufacturer's recommendations.

**Measurement of gene expression by RT-qPCR**

*L. interrogans* WT, *sodB*-expressing, *katG*-expressing and *sodB*-*katG*-expressing strains were cultivated in EMJH medium until exponential phase. Harvested bacteria were resuspended in 1 ml TRIzol (ThermoFisher Scientific) and stored at -80°C. Nucleic Acids were extracted with chloroform and precipitated with isopropanol as described before[23]. Contaminating genomic DNA was removed by DNAse treatment using the RNAse-free Turbo DNA-free turbo kit (ThermoFisher Scientific) as described by the manufacturer. cDNA synthesis was performed with the cDNA synthesis kit (Biorad) according to the manufacturer's recommendation. Quantitative PCR was conducted with the SsoFast EvaGreen Supermix (Biorad) as previously described[22]. Gene expression was measured with primers described in Table S11 using *flaB* (LIMLP_09410) as a reference gene.

**Determination of bacteria viability**

Leptospires survival was determined by incubating exponentially growing *L. interrogans* (≈ $10^8$/ml) in EMJH in the presence or absence of 10 or 50 µM paraquat for 1 h. Colony-forming unit quantification was performed by diluting bacteria in EMJH and plating on solid EMJH medium. After one month incubation at 30°C, colonies were counted and percent survival (%

of colony forming unit, CFU) was calculated as the ratio of CFU for bacteria incubated in the presence of paraquat to that for bacteria incubated in the absence of paraquat.

**Infection experiments**

*L. interrogans* WT and *sodB*-expressing strains were cultivated in EMJH medium until the exponential phase and counted under a dark-field microscope using a Petroff-Hauser cell. $10^6$ leptospires (in 0.5 ml) were injected intraperitoneally in groups of 4 male 4 weeks-old Syrian Golden hamsters (RjHan:AURA, Janvier Labs). Animals were monitored daily and sacrificed by $CO_2$ inhalation when endpoint criteria were met (sign of distress, morbidity).

**Ethics Statement**

The protocol for animal experimentation was reviewed by the Institut Pasteur (Paris, France), the competent authority, for compliance with the French and European regulations on Animal Welfare and with Public Health Service recommendations. This project has been reviewed and approved (CETEA 220016) in accordance with the recommendations of the Specific Guide for the Care and the Use of Laboratory Animals of the Institut Pasteur, according to the European Directive (2010/63/UE) and the corresponding French law on animal experimentation.

**Hydrogen sulfide production**

45 ml of WT exponentially growing *L. interrogans* were exposed or not to 50 µM or 200 µM paraquat for 1.5 hours. Cultures were then harvested by centrifugation and resuspended in 1 ml of PBS. 300 µl of each suspension was used to determine hydrogen sulfide production using the OxiSelect Free Hydrogen Sulfide Gas Assay Kit (Xan-5084, Cell Biolabs) following the manufacturer's recommendations with one hour incubation. A positive control for $H_2S$ production was included using 200 µM $Na_2S$ in water.

**Aconitase activity**

Aconitase activity were assayed in the total extracts of *Leptospira* (2.5 µg of total extract) using the colorimetric Aconitase Activity Assay Kit (MAK051, Sigma-Aldrich) following the manufacturer's protocol without activation solution. Isocitrate production was calibrated using a standard curve from 0-20 nmoles. Activity was determined in mU/ml and further normalized to the untreated control.

**Determination of amino acid abundance**

Amino acid abundance in four replicates were obtained by counting the number of each amino acid per peptide and multiplying by the number of spectra associated to each peptide. These values were then summed per amino acid and condition and normalized by the total amount of amino acids per condition to obtain the frequency of each amino acid. Some representations further normalized each percentage by the untreated replicates in a paired manner.

### SUPPLEMENTARY FIGURES

**Figure S1**: Phylogenomic analysis and distribution of SOSE in the *Spirochaetes* phylum.

**Figure S2**: *SodB* and *SodA* loci in *Leptospira* spp.

**Figure S3**: Expression of *L. biflexa sodB* and *katG* in *L. interrogans*.

**Figure S4**: Adaptation of pathogenic *Leptospira* spp. to superoxide.

**Figure S5**: Description of the locus LIMLP_14415-14460.

**Figure S6**: RNASeq analysis in the presence of paraquat.

**Figure S7**: Metabolic pathways upregulated with superoxide.

**Figure S8**: LeuA1 and LeuA2 *of Leptospira* spp.

**Figure S9**: Amino acid abundance.

**Figure S10**: Amino acid abundance upon exposure to superoxide.

**Figure S11**: Effect of exposure to superoxide on amino acid oxidation.

**Figure S12**: Role of sulfate assimilatory reduction on *L. interrogans* fitness with superoxide.

### SUPPLEMENTARY TABLES

A description of the content is available within each table

**Table S1.** Distribution of SOD and SOR in bacterial pathogenic species.

**Table S2.** Species of the *Spirochaetes* phylum.

**Table S3**: Mutations identified in the superoxide-adapted strain.

**Table S4**: Complete data set of RNASeq of the superoxide-adapted strain.

**Table S5**: Complete data set of the RNASeq of *L. interrogans* upon exposure to 10 and 50 µM paraquat.

**Table S6**: Upregulated genes upon exposure to superoxide.

**Table S7**: Downregulated genes upon exposure to superoxide.

**Table S8**: Proteome of *L. interrogans* upon exposure to 10 and 50 µM paraquat

**Table S9**: Comparison of DEGs of *L. interrogans* upon exposure to 50 µM paraquat and 1 mM $H_2O_2$.

**Table S10**: Plasmids and strains used in this study.

**Table S11**: Primers used in this study.

### SUPPLEMENTARY FIGURE LEGENDS

**Figure S1. Phylogenomic analysis and distribution of SOSE in the *Spirochaetes* phylum.**

(A) Phylogenetic tree of SOSEs (SOR or SOD) in the *Spirochaetes* phylum.

(B) Phylogenetic distribution of SOSE (SOD or SOR) in the *Spirochaetes* phylum. *Treponemataceae* are in blue, *Spirochaetecales* are in light blue, *Borreliaceae* are in red, *Brachyspirales* are in yellow, and *Leptospirales* are in cyan. The presence or absence of a SOSE are indicated in blue and grey, respectively. The aerobicity is also indicated.

**Figure S2. *SodB* and *SodA* loci in *Leptospira* spp.**

The *sodB* locus of S1 and S2 clades and the corresponding loci in P1 and P2 clades are represented. The *sodA* locus of *L. perolatii* is also represented.

**Figure S3. Expression of *L. biflexa sodB* and *katG* in *L. interrogans*.**

(A-B) *SodB* (A) and *katG* (B) expressions were measured in the WT (circle), *sod*-expressing (square), *katG*-expressing (triangle) and *sod*- and *katG*-expressing (inverted triangle) *L. interrogans* strains by RT-qPCR. Data are mean and SD of three independent experiments.

(C) The virulence of the WT (black circle) or *sod*-expressing (green square) was evaluated in Syrian golden hamsters (n=4) using $10^6$ leptospires injected by peritoneal route.

(D) SOD activity was measured in the WT *L. interrogans* strain untreated (grey bar) or exposed to 50 μM paraquat (PQ) (blue bar), and in *Escherichia coli* C600 strain (used as a positive control) (red bar).

**Figure S4. Adaptation of pathogenic *Leptospira* spp. to superoxide.**

(A) Schematic representation of the pipeline for experimental adaptation of *L. interrogans* to superoxide. The adaptation was performed by cultivating *L. interrogans* in the presence of 2 μM paraquat (PQ). The cycle numbers are the number of *in vitro* passages.

(B) Growth curve of WT strain at cycle 1 (upper panel) or 2 (lower panel) in the absence (black symbols) or presence (red symbols) of 2 μM paraquat (PQ). Growth was assessed by measure of absorbance at 420 nm. Data are mean and SD of four independent experiments.

(C) Growth curve of non-adapted (control) and adapted strains at cycle 7 or 46 (as indicated) in the absence (black circle) or presence (red square) of 2 μM paraquat (PQ). Growth was assessed as described in (B). Data are mean and SD of three independent experiments.

(D) The time to exit lag phase (OD>0.05) was plotted at cycle 1, 2 and for the control and adapted strain at cycle 7 and 46 (as indicated) in the absence (white bar, circle) or presence (red bar, square) of 2 μM PQ.

(E) Growth curve of non-adapted (control, black circle) and adapted (green square) strains in the presence of 0.5 mM $H_2O_2$. Growth was assessed as described in (B). Data are mean and SD of three independent experiments.

**Figure S5. Description of the locus LIMLP_14415-14460.**

(A) Locus LIMLP_14415-14460 composition. Annotations are according to the Microbial Genome Annotation & Analysis Platform (MicroScope, *L. interrogans* serovar Manilae strain UP-MMC-NIID LP).

(B) Schematic representation of the locus. Each factor is represented with its predicted structure (determined by Alphafold and available in Uniprot) and putative cellular localization (as predicted by PsortB). The $Log_2FC$ of their expression in the adapted strain compared to the control strain are also indicated.

**Figure S6. RNASeq analysis in the presence of paraquat.**

(A) Survival of *L. interrogans* WT strain upon exposure for 1h to 0 (circle, grey bar), 10 (square, green bar), and 50 μM (triangle, blue bar) paraquat (PQ). Survival has been determined by the percentage of colony-forming unit (CFU) and normalized by that of untreated control. Data are mean and SD of three independent experiments. *: p-value<0.02.

(B) Gene ontology analysis of enriched pathway in upregulated (right) and downregulated (left) DEGs upon exposure to paraquat.

**Figure S7. Metabolic pathways upregulated with superoxide**

Leucine biosynthesis (A) and sulfate assimilatory reduction (B) pathways are schematically represented. For each reaction, gene names are indicated (when identified). Differential expression (expressed as $Log_2FC$) is indicated into parenthesis for each factor of the leucine biosynthesis pathway.

**Figure S8. LeuA1 and LeuA2 of *Leptospira* spp.**

(A) Sequence alignment of LeuA1 (LIMLP_08570) and LeuA2 (LIMLP_15720) in *Leptospira* genus. The schematic domain organization of a typical LeuA is represented at the top. Conserved residues are highlighted in blue.

(B) Representativity of leucine residues in *L. interrogan*s WT strain proteome upon exposure to 0 (grey bar, circle) 10 (green bar, square) and 50 (blue bar, triangle) $\mu$M paraquat. Data are mean and SD of four independent experiments. The predicted theoretical value (10.41%) is indicated by the dashed line.

**Figure S9. Amino acid abundance.**

Percentage of each amino acid (indicated by the letter code) in the genome of *L. interrogans* as determined *in silico*. Boxplots represent the median and first and third quartiles, and the blue dots in the background represent the percentage of each amino acid in each individual protein.

**Figure S10. Amino acid abundance upon exposure to superoxide.**

Percentage of each amino acid in the expressed proteome of *L. interrogans* as determined by peptide analysis through mass spectrometry. Amino acids are designated by letter code, oxidized Met is indicated as B, bi-oxidized Cys as J, and tri-oxidized Cys as O. Data are boxplots of the median and first and third quartiles with the four individual replicates plotted on top per condition. The Y axis is logarithmic.

**Figure S11. Effect of exposure to superoxide on amino acid oxidation.**

(A) Cysteine oxidation was quantified by mass spectrometry in *L. interrogans* WT strain exposed 1h to 10 and 50 µM paraquat (as indicated). The X axis includes all proteins with peptides found in all replicates (*N*=1434) and ordered by localization in the genome. Each peak represents a minimal boxplot with median and first and third quartiles. DnaG is the protein with the highest oxidation ratio upon superoxide exposure (indicated by an asterisk). The median value of the oxidation ratio (and its range) of DnaG is indicated.

(B) Growth (upper panel) and genome replication (lower panel) of *L. interrogans* WT strain when 50 µM paraquat is added in exponential phase (48 hours post-inoculation, blue line, triangle). The black line (circle) represents untreated samples. Data are mean and SD of three independent biological replicates.

(C) Representativity of cysteine residues in the up- (in red) and downregulated (in blue) peptides (left panel) and genes (right panel) upon exposure to 50 µM paraquat.

(D) Aconitase activity in a total extract of *L. interrogans* WT strain upon exposure to 0 (grey bar, circle) 10 (green bar, square) and 50 (blue bar, triangle) µM paraquat. Data is geometric mean and geometric SD of three independent biological replicates.

(E) Expression values (expressed as $Log_2FC$) of Fe-S containing proteins (as predicted by Metal Predator[24]) upon exposure to 10 and 50 µM paraquat.

**Figure S12. Role of sulfate assimilatory reduction on *L. interrogans* fitness with superoxide.**

(A-D) Growth of *L. interrogans* WT strain in the absence or presence of 3 µM paraquat (PQ) in a EMJH medium complemented with 200 µM $Na_2SO_3^{2-}$ (A), 0.1 mM $Na_2S$ or 1 mM NaSH (B),

200 μM cysteine (C) 1 mM cystine dissolved in Tris-HCl (D). Growth was assessed as described in Figure S4. Data are mean and SD of three independent experiments.

(E) $H_2S$ production was determined in *L. interrogans* WT strain exposed for 1h without or with 50 and 200 μM paraquat (as indicated). Data mean and SD of three independent experiments.

(F) Growth of *L. interrogans* WT (black circle) and *metY* mutant (Man 513) (red square) strains in the absence (solid line) or presence of 3 μM paraquat (dashed line). Data are mean and SD of three independent experiments.

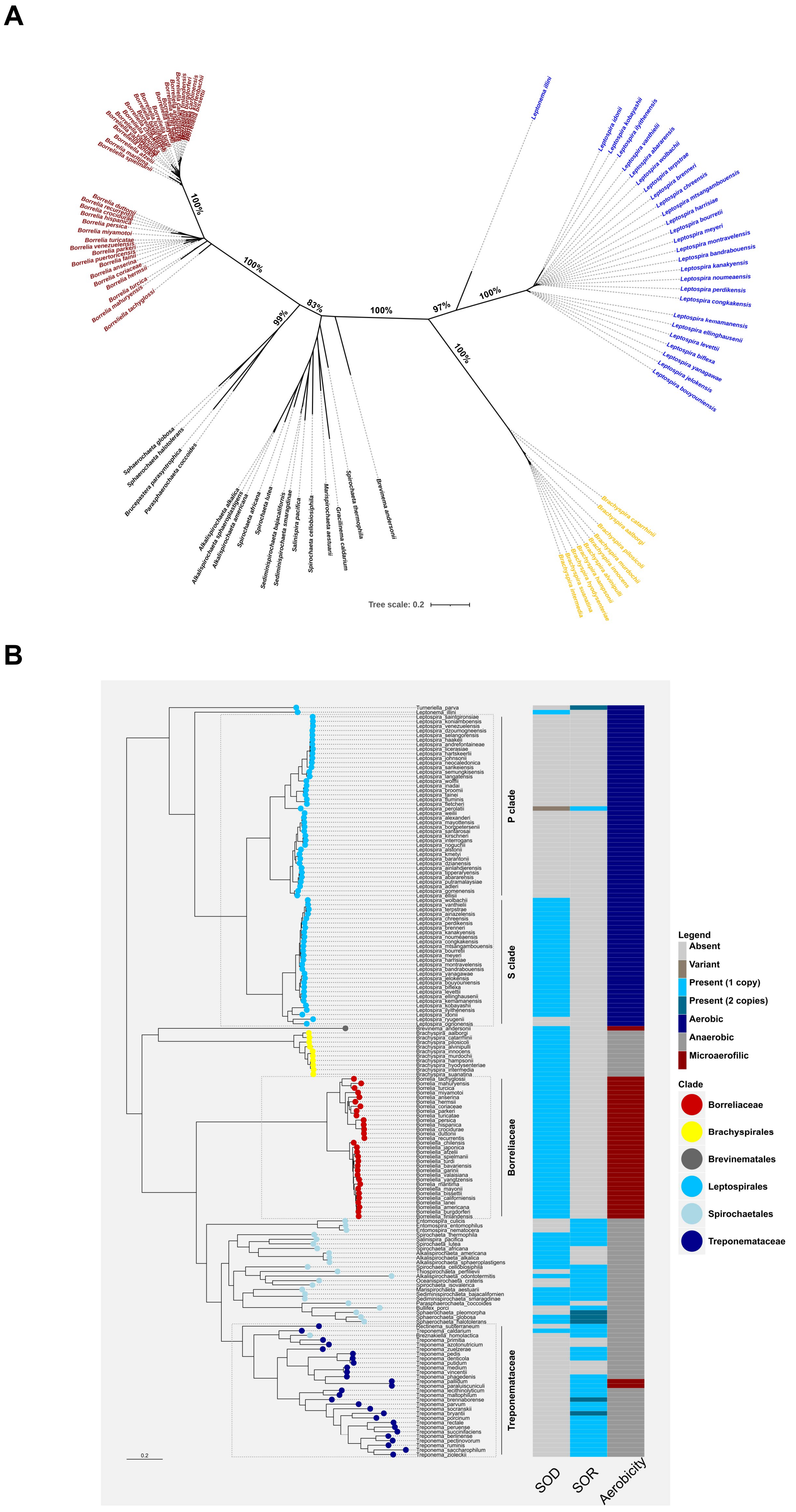
A
100%
100%
99%
83%
100%
97%
100%
100%
Tree scale: 0.2
B
P clade
S clade
Borreliaceae
Treponemataceae
0.2
SOD
SOR
Aerobicity
Legend
Absent
Variant
Present (1 copy)
Present (2 copies)
Aerobic
Anaerobic
Microaerofilic
Clade
Borreliaceae
Brachyspirales
Brevinematales
Leptospirales
Spirochaetales
Treponemataceae

Figure S1

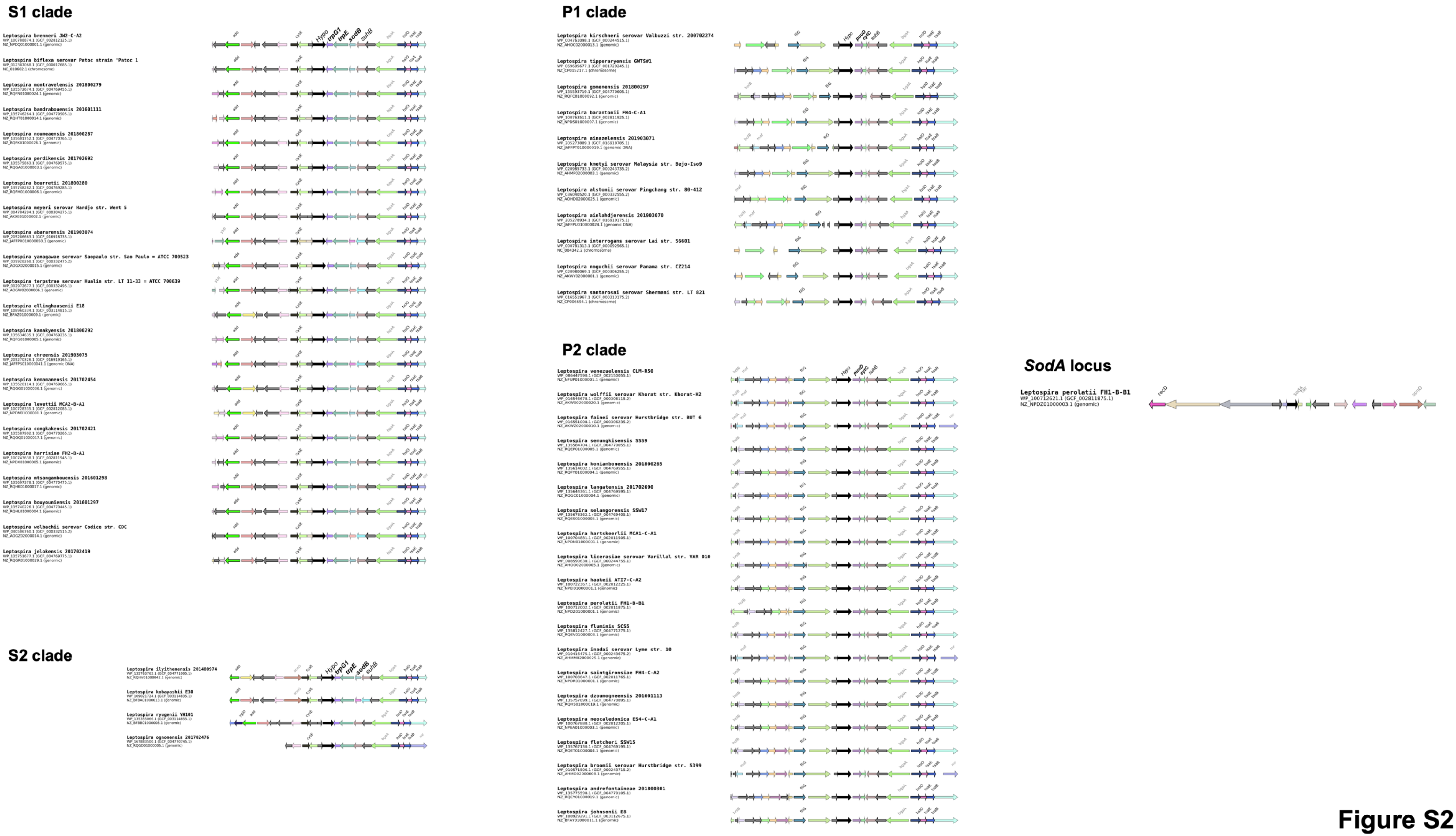
S1 clade
Leptospira brenneri JW2-C-A2
Leptospira biflexa serovar Patoc strain 'Patoc 1
Leptospira montravelensis 201800279
Leptospira bandrabouensis 201601111
Leptospira noumeaensis 201800287
Leptospira perdikensis 201702692
Leptospira bourretii 201800280
Leptospira meyeri serovar Hardjo str. Went 5
Leptospira abararensis 201903074
Leptospira yanagawae serovar Saopaulo str. Sao Paulo = ATCC 700523
Leptospira terpstrae serovar Hualin str. LT 11-33 = ATCC 700639
Leptospira ellinghausenii E18
Leptospira kanakyensis 201800292
Leptospira chreensis 201903075
Leptospira kemamanensis 201702454
Leptospira levettii MCA2-B-A1
Leptospira congkakensis 201702421
Leptospira harrisiae FH2-B-A1
Leptospira mtsangambouensis 201601298
Leptospira bouyouniensis 201601297
Leptospira wolbachii serovar Codice str. CDC
Leptospira jelokensis 201702419
S2 clade
Leptospira ilyithenensis 201400974
Leptospira kobayashii E30
Leptospira ryugenii YH101
Leptospira ognonensis 201702476
P1 clade
Leptospira kirschneri serovar Valbuzzi str. 200702274
Leptospira tipperaryensis GWTS#1
Leptospira gomenensis 201800297
Leptospira barantonii FH4-C-A1
Leptospira ainazelensis 201903071
Leptospira kmetyi serovar Malaysia str. Bejo-Iso9
Leptospira alstonii serovar Pingchang str. 80-412
Leptospira ainlahdjerensis 201903070
Leptospira interrogans serovar Lai str. 56601
Leptospira noguchii serovar Panama str. CZ214
Leptospira santarosai serovar Shermani str. LT 821
P2 clade
Leptospira venezuelensis CLM-R50
Leptospira wolffii serovar Khorat str. Khorat-H2
Leptospira fainei serovar Hurstbridge str. BUT 6
Leptospira semungkisensis SSS9
Leptospira koniambonensis 201800265
Leptospira langatensis 201702690
Leptospira selangorensis SSW17
Leptospira hartskeerlii MCA1-C-A1
Leptospira licerasiae serovar Varillal str. VAR 010
Leptospira haakeii ATI7-C-A2
Leptospira perolatii FH1-B-B1
Leptospira fluminis SCS5
Leptospira inadai serovar Lyme str. 10
Leptospira saintgironsiae FH4-C-A2
Leptospira dzoumogneensis 201601113
Leptospira neocaledonica ES4-C-A1
Leptospira fletcheri SSW15
Leptospira broomii serovar Hurstbridge str. 5399
Leptospira andrefontaineae 201800301
Leptospira johnsonii E8
SodA locus
Leptospira perolatii FH1-B-B1

Figure S2

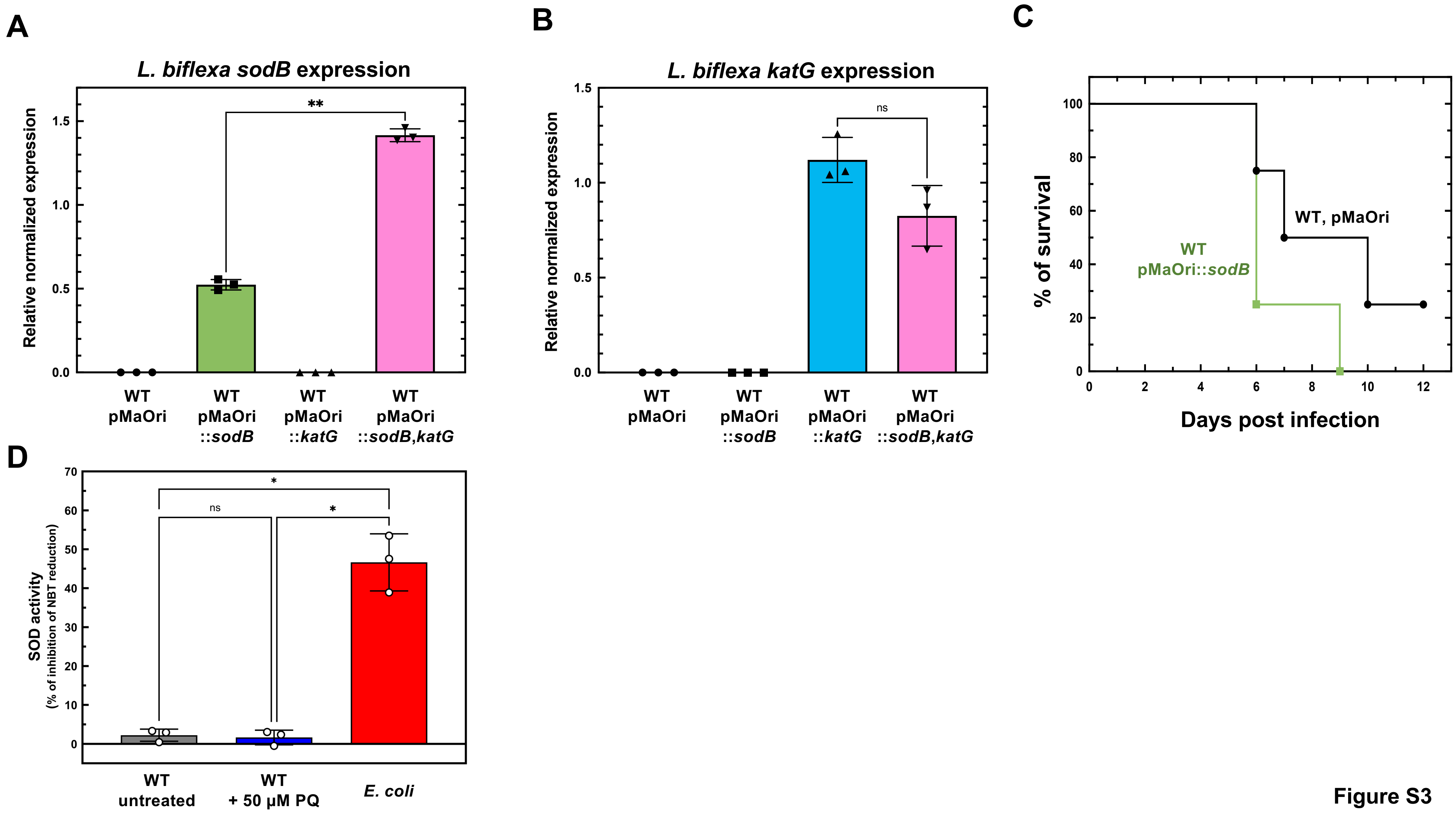
A
L. biflexa sodB expression
Relative normalized expression
0.0
0.5
1.0
1.5
**
WT pMaOri
WT pMaOri ::sodB
WT pMaOri ::katG
WT pMaOri ::sodB,katG
B
L. biflexa katG expression
Relative normalized expression
0.0
0.5
1.0
1.5
ns
WT pMaOri
WT pMaOri ::sodB
WT pMaOri ::katG
WT pMaOri ::sodB,katG
C
% of survival
0
20
40
60
80
100
WT, pMaOri
WT pMaOri::sodB
0
2
4
6
8
10
12
Days post infection
D
SOD activity
(% of inhibition of NBT reduction)
0
10
20
30
40
50
60
70
*
ns
*
WT untreated
WT + 50 µM PQ
E. coli

Figure S3

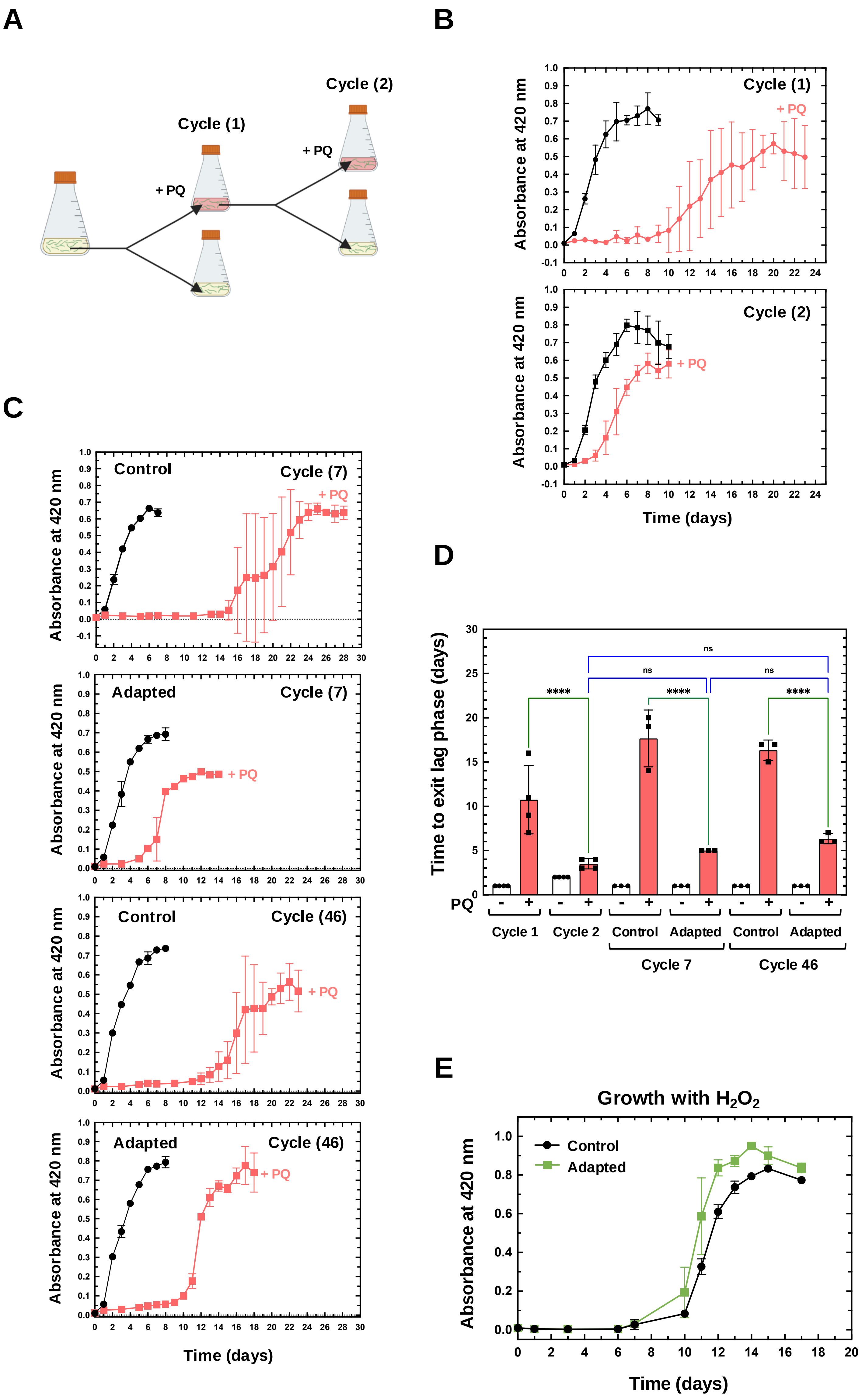
A
Cycle (1)
Cycle (2)
+ PQ
+ PQ
B
Absorbance at 420 nm
Cycle (1)
+ PQ
Cycle (2)
+ PQ
Time (days)
C
Absorbance at 420 nm
Control
Cycle (7)
+ PQ
Adapted
Cycle (7)
+ PQ
Control
Cycle (46)
+ PQ
Adapted
Cycle (46)
+ PQ
Time (days)
D
Time to exit lag phase (days)
ns
ns
ns
****
****
****
PQ
- + - + - + - + - + - +
Cycle 1
Cycle 2
Control
Adapted
Control
Adapted
Cycle 7
Cycle 46
E
Growth with $H_2O_2$
Control
Adapted
Absorbance at 420 nm
Time (days)

**Figure S4**

**A**

| Locus | | Annotation |
|---|---|---|
| LIMLP_14415 | LIMLP_RS14380 | ArsR/SmtB Transcriptional Regulator |
| LIMLP_14420 | LIMLP_RS14385 | DoxX protein family |
| LIMLP_14425 | LIMLP_RS14390 | START domain-containing protein |
| LIMLP_14430 | LIMLP_RS14395 | START domain-containing protein |
| LIMLP_14435 | LIMLP_RS14400 | START domain-containing protein |
| LIMLP_14440 | LIMLP_RS14405 | START domain-containing protein |
| LIMLP_14445 | LIMLP_RS14410 | Hypothetical |
| LIMLP_14450 | LIMLP_RS14415 | Hypothetical |
| LIMLP_14455 | LIMLP_RS14420 | DHFR domain-containing protein |
| LIMLP_14460 | LIMLP_RS14425 | MFS transporter |

**B**

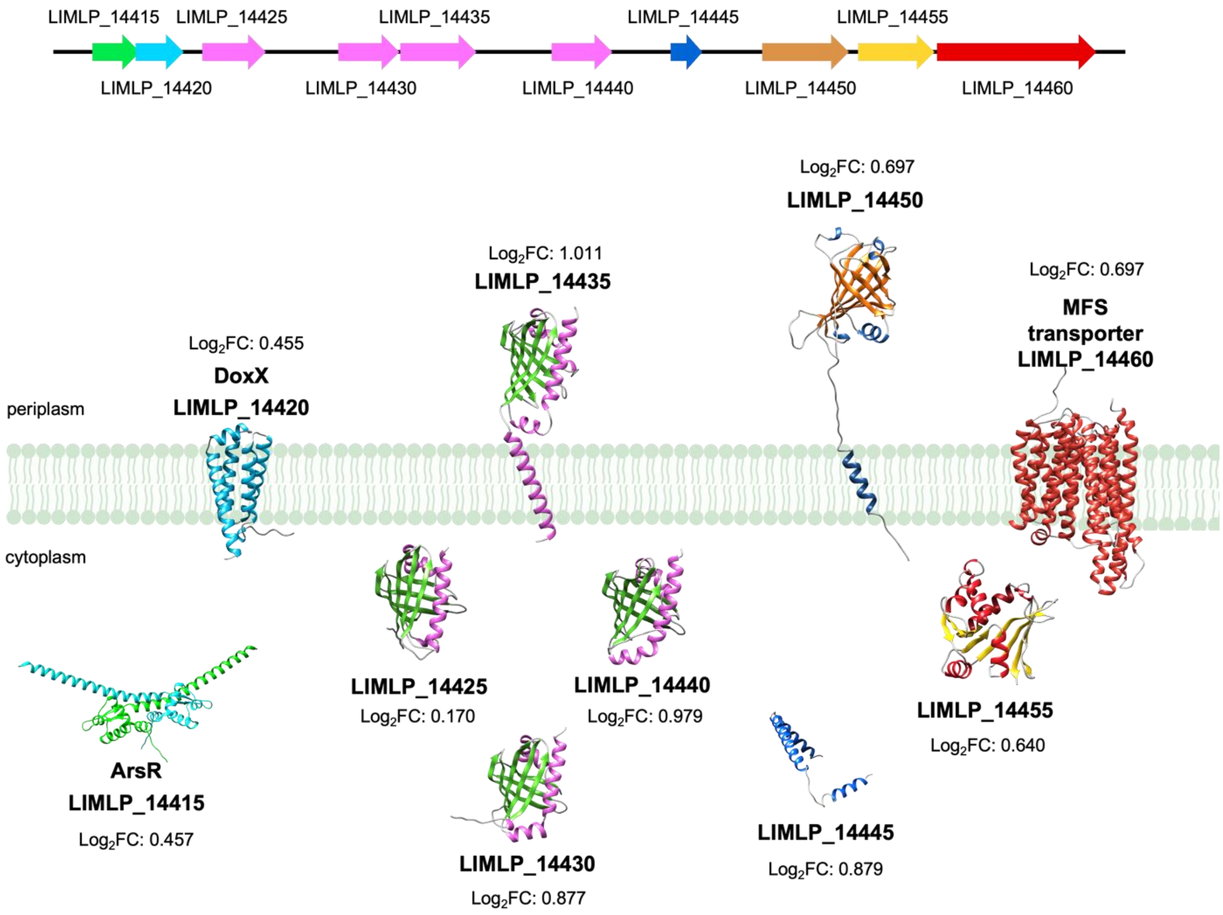


**Figure S5**

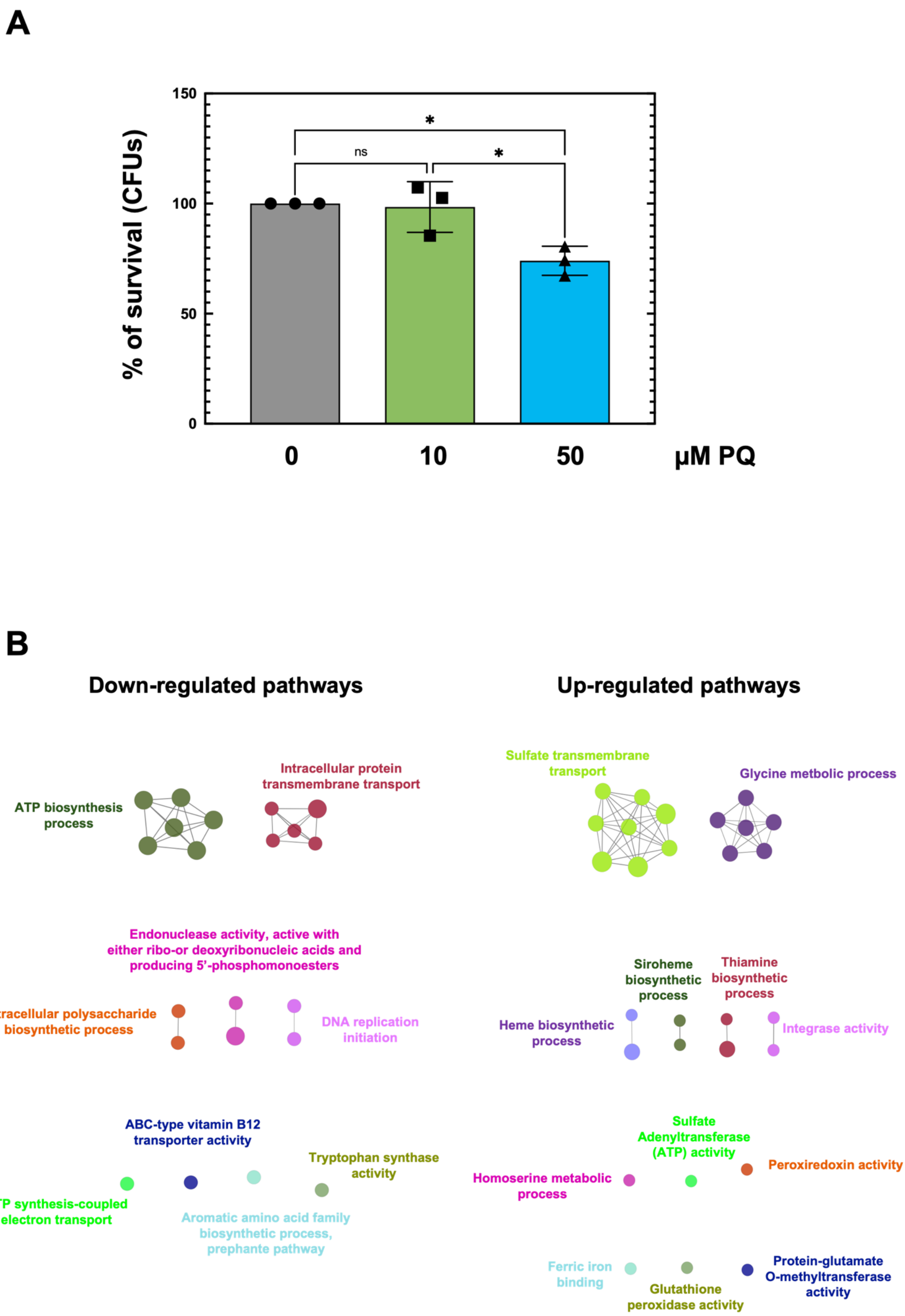

A
150
100
50
0
% of survival (CFUs)
*
ns
*
0
10
50
µM PQ
B
Down-regulated pathways
Up-regulated pathways
ATP biosynthesis process
Intracellular protein transmembrane transport
Endonuclease activity, active with either ribo-or deoxyribonucleic acids and producing 5'-phosphomonoesters
Extracellular polysaccharide biosynthetic process
DNA replication initiation
ABC-type vitamin B12 transporter activity
Tryptophan synthase activity
ATP synthesis-coupled electron transport
Aromatic amino acid family biosynthetic process, prephante pathway
Sulfate transmembrane transport
Glycine metbolic process
Siroheme biosynthetic process
Thiamine biosynthetic process
Heme biosynthetic process
Integrase activity
Sulfate Adenyltransferase (ATP) activity
Homoserine metabolic process
Peroxiredoxin activity
Ferric iron binding
Glutathione peroxidase activity
Protein-glutamate O-methyltransferase activity


Figure S6

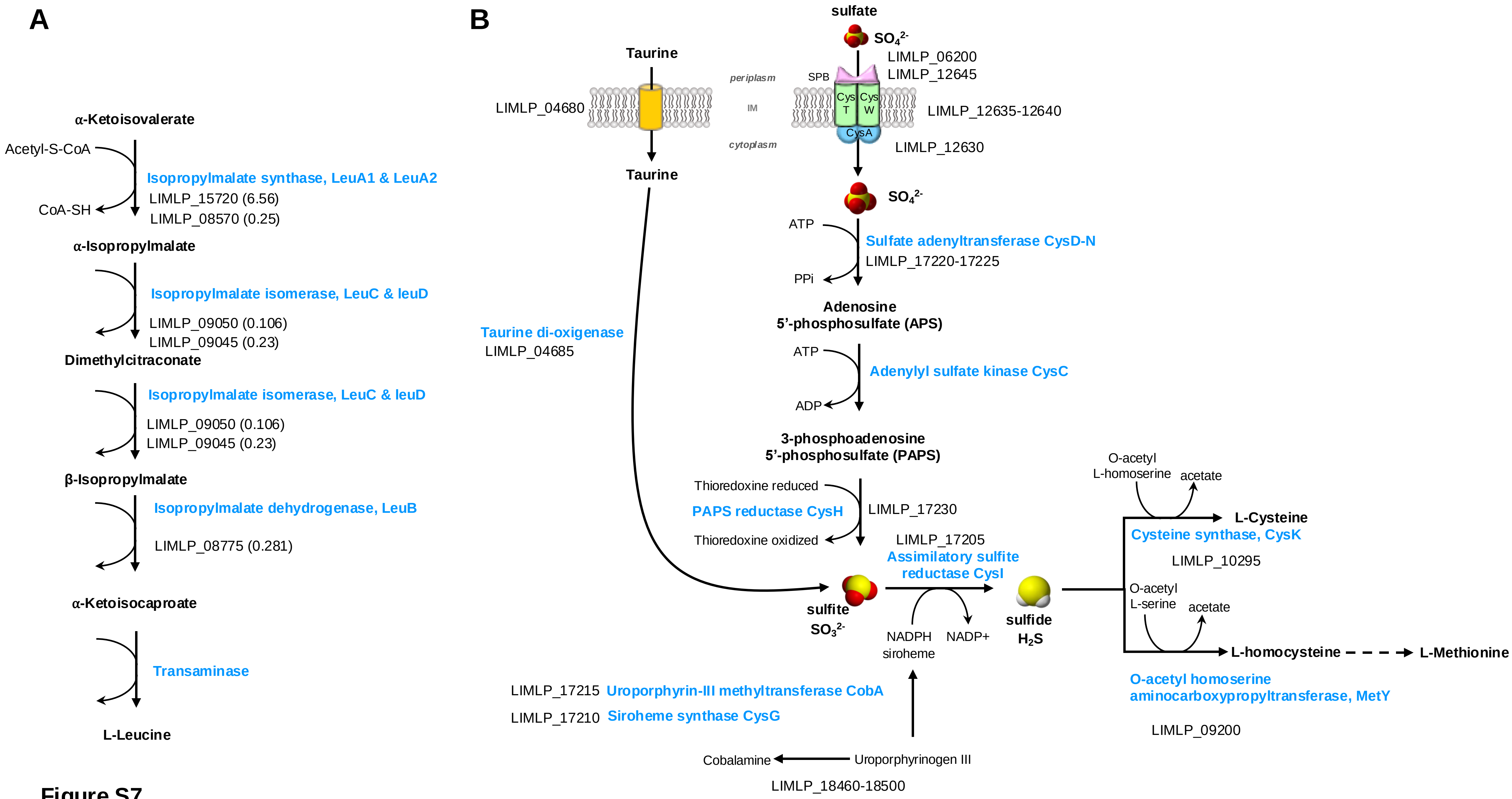
A
α-Ketoisovalerate
Acetyl-S-CoA
CoA-SH
Isopropylmalate synthase, LeuA1 & LeuA2
LIMLP_15720 (6.56)
LIMLP_08570 (0.25)
α-Isopropylmalate
Isopropylmalate isomerase, LeuC & leuD
LIMLP_09050 (0.106)
LIMLP_09045 (0.23)
Dimethylcitraconate
Isopropylmalate isomerase, LeuC & leuD
LIMLP_09050 (0.106)
LIMLP_09045 (0.23)
β-Isopropylmalate
Isopropylmalate dehydrogenase, LeuB
LIMLP_08775 (0.281)
α-Ketoisocaproate
Transaminase
L-Leucine
B
Taurine
LIMLP_04680
periplasm
IM
cytoplasm
Taurine
Taurine di-oxigenase
LIMLP_04685
sulfate
$SO_4^{2-}$
LIMLP_06200
LIMLP_12645
SPB
Cys T
Cys W
CysA
LIMLP_12635-12640
LIMLP_12630
$SO_4^{2-}$
ATP
PPi
Sulfate adenyltransferase CysD-N
LIMLP_17220-17225
Adenosine 5'-phosphosulfate (APS)
ATP
ADP
Adenylyl sulfate kinase CysC
3-phosphoadenosine 5'-phosphosulfate (PAPS)
Thioredoxine reduced
PAPS reductase CysH
Thioredoxine oxidized
LIMLP_17230
LIMLP_17205
Assimilatory sulfite reductase CysI
sulfite
$SO_3^{2-}$
NADPH
NADP+
siroheme
sulfide
$H_2S$
O-acetyl L-homoserine
acetate
L-Cysteine
Cysteine synthase, CysK
LIMLP_10295
O-acetyl L-serine
acetate
L-homocysteine
L-Methionine
O-acetyl homoserine aminocarboxypropyltransferase, MetY
LIMLP_09200
LIMLP_17215 Uroporphyrin-III methyltransferase CobA
LIMLP_17210 Siroheme synthase CysG
Cobalamine
Uroporphyrinogen III
LIMLP_18460-18500

**Figure S7**

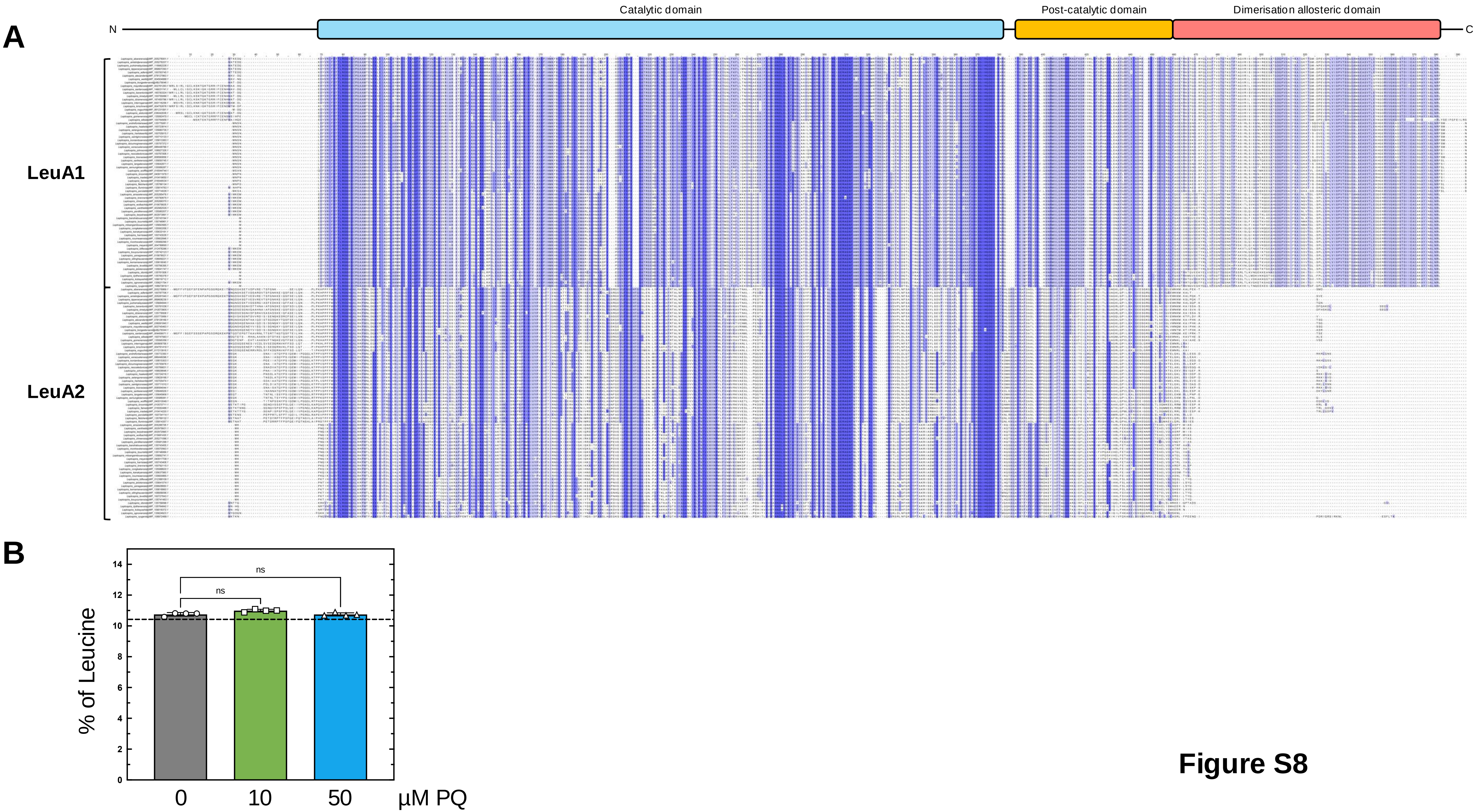
A
Catalytic domain
Post-catalytic domain
Dimerisation allosteric domain
N
C
LeuA1
LeuA2
B
% of Leucine
14
12
10
8
6
4
2
0
ns
ns
0
10
50
µM PQ

**Figure S8**

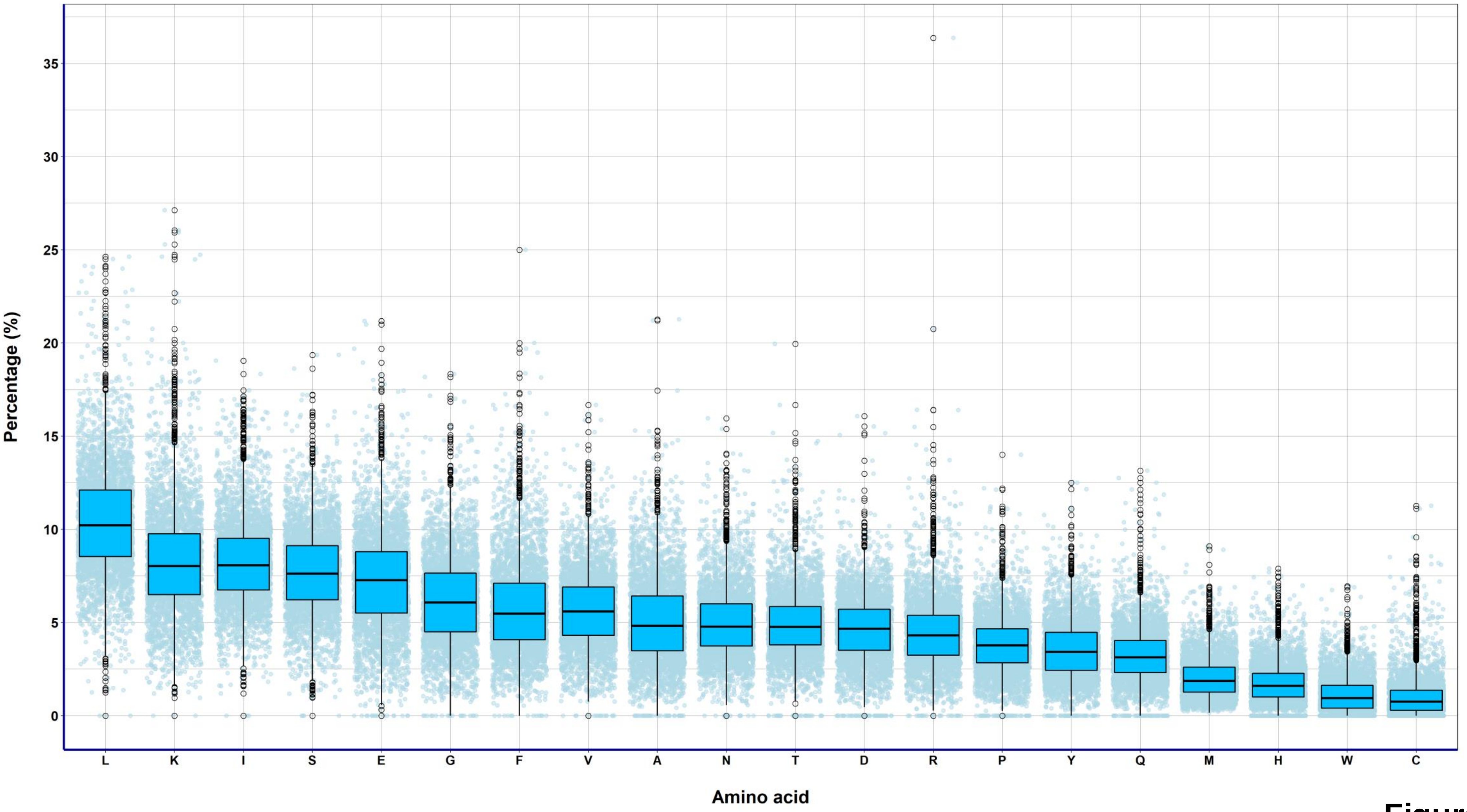
Percentage (%)
0
5
10
15
20
25
30
35
Amino acid
L
K
I
S
E
G
F
V
A
N
T
D
R
P
Y
Q
M
H
W
C

**Figure S9**

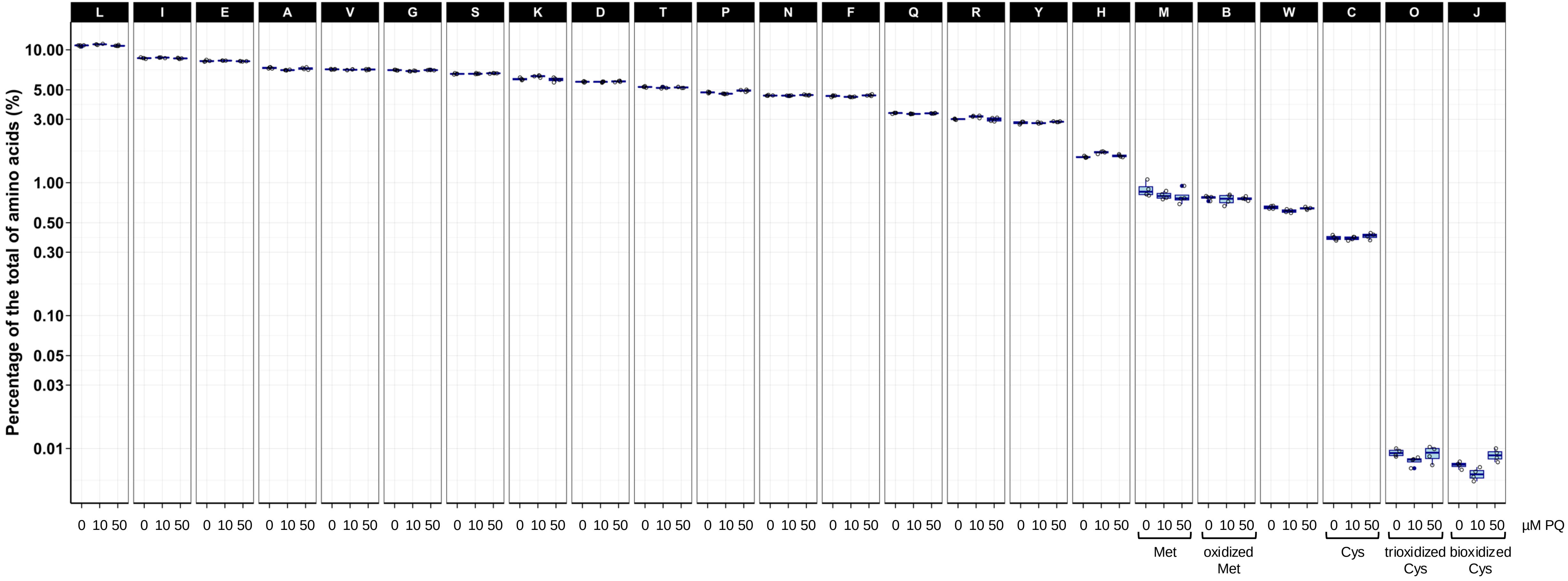

L
I
E
A
V
G
S
K
D
T
P
N
F
Q
R
Y
H
M
B
W
C
O
J
Percentage of the total of amino acids (%)
10.00
5.00
3.00
1.00
0.50
0.30
0.10
0.05
0.03
0.01
0 10 50
µM PQ
Met
oxidized Met
Cys
trioxidized Cys
bioxidized Cys


Figure S10

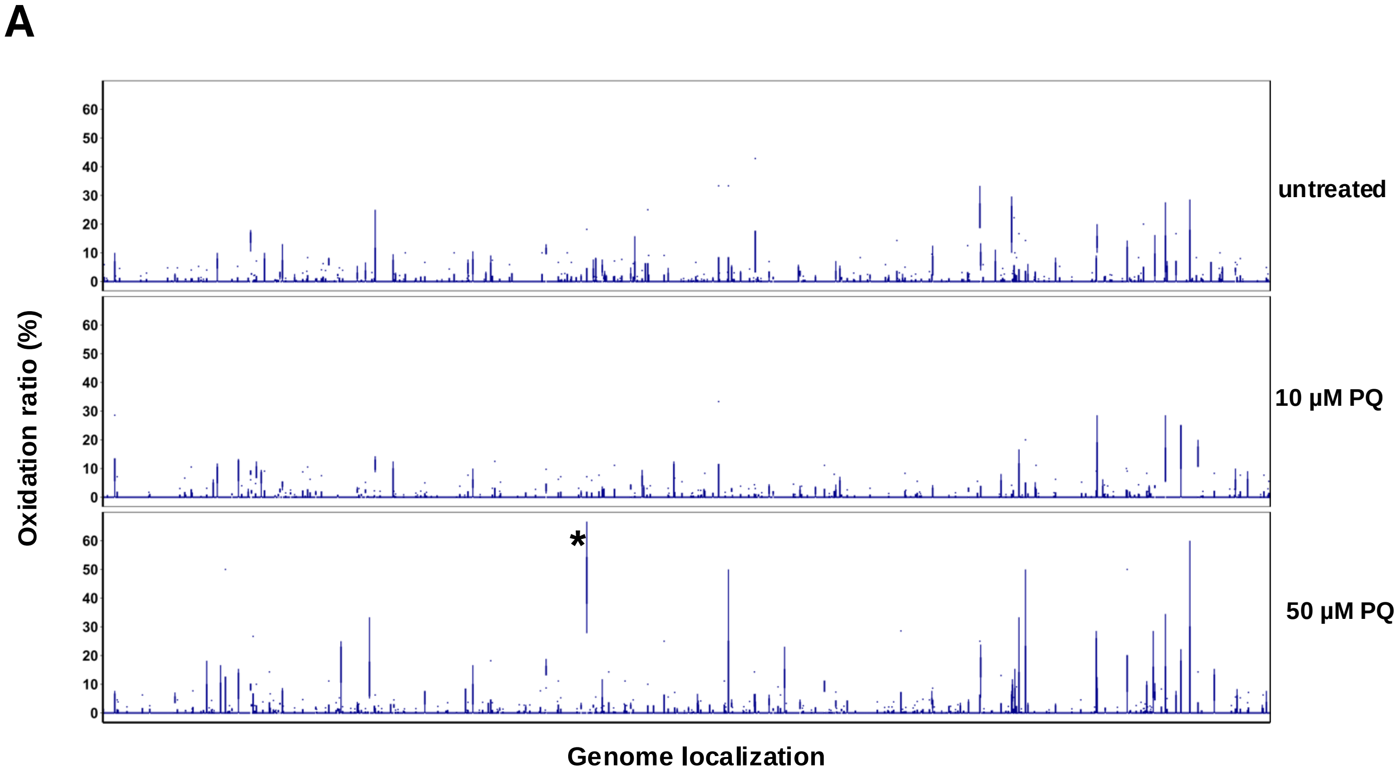


| ORF Id | Annotation | Pathway | Oxidation ratio |
|---|---|---|---|
| LIMLP_08450 | DnaG primase | DNA replication | 45.8% (27.8-66.7) |

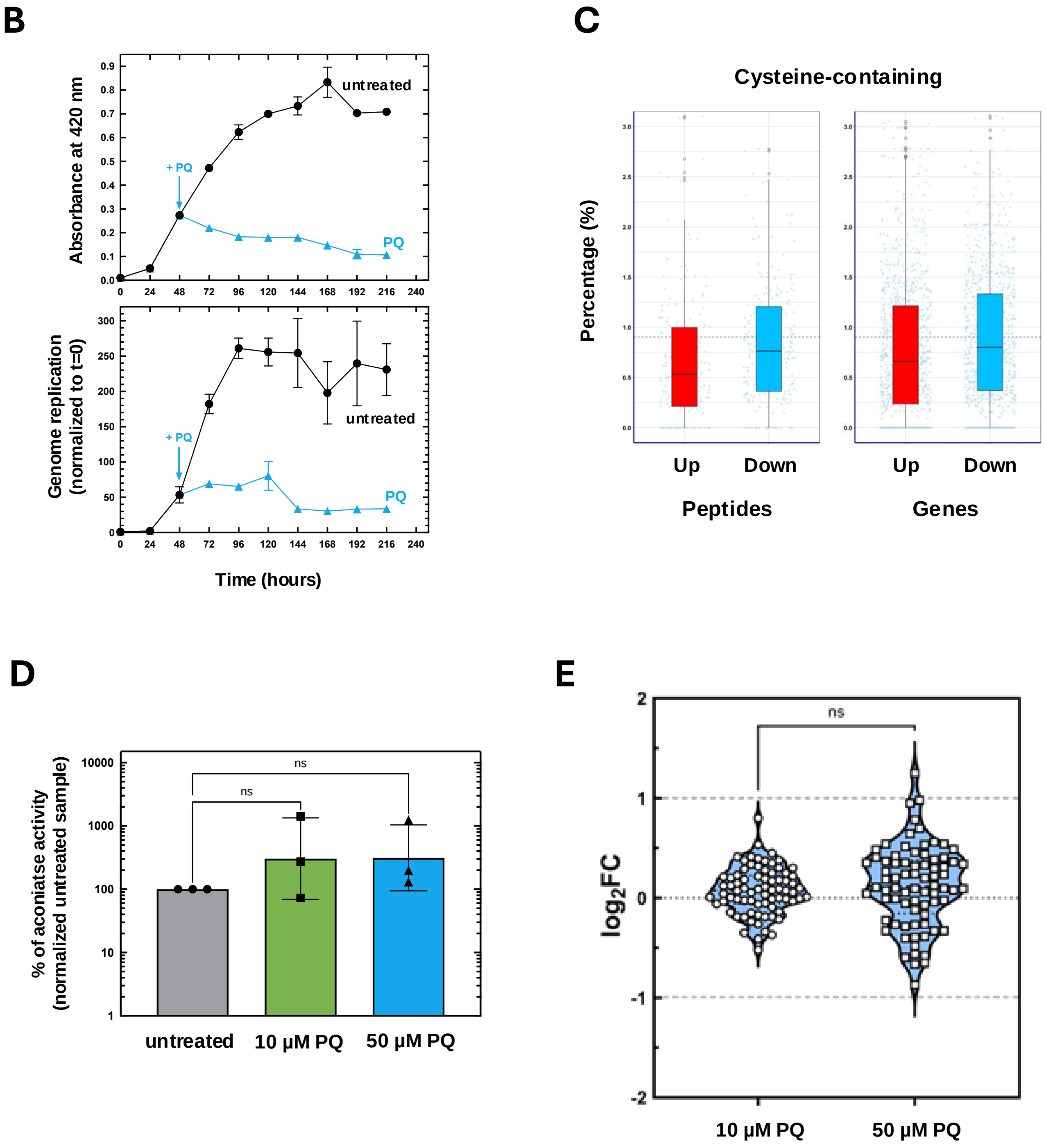


Figure S11

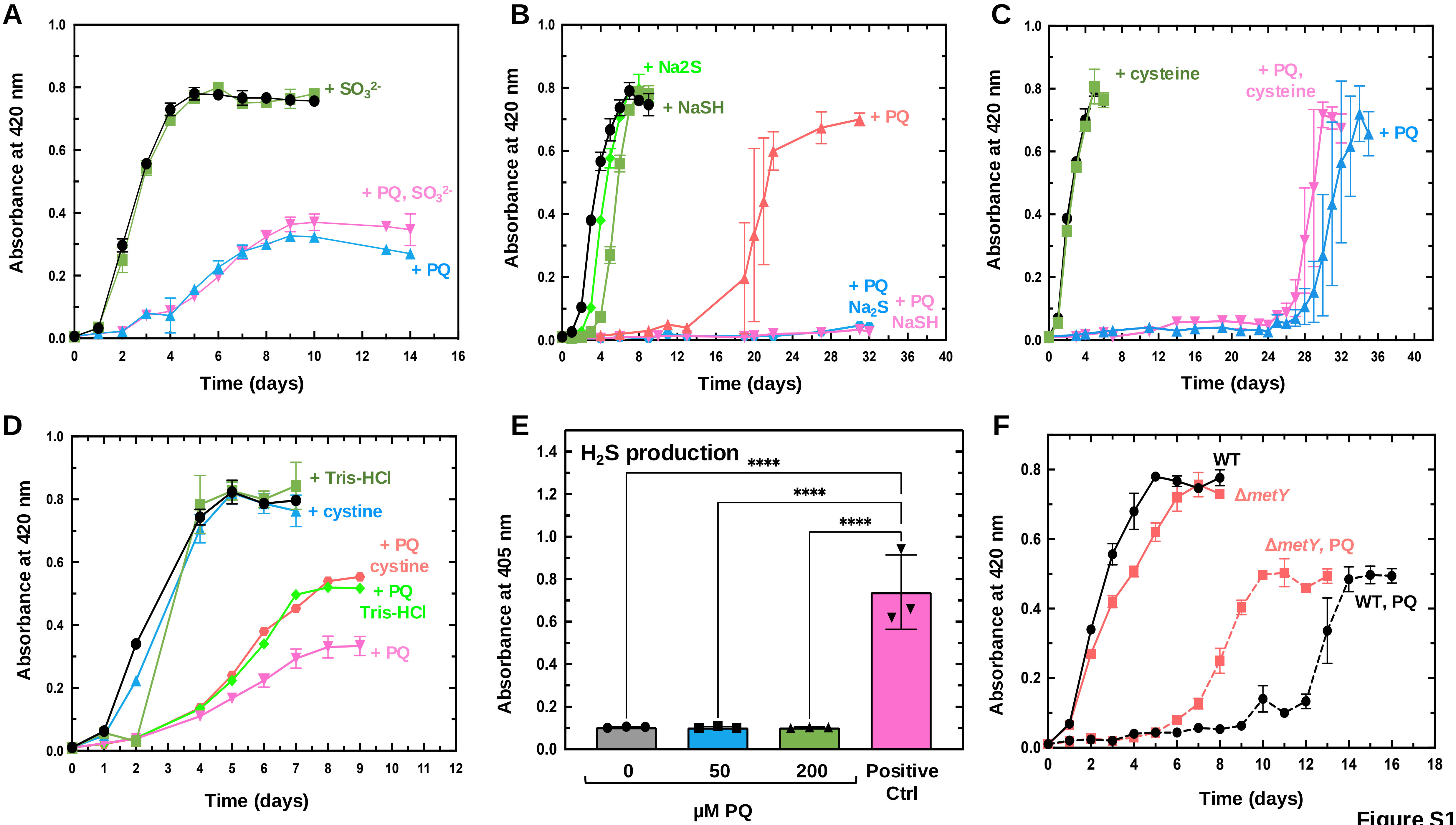
A
B
C
D
E
F
Absorbance at 420 nm
Time (days)
+ SO$_3^{2-}$
+ PQ, SO$_3^{2-}$
+ PQ
+ Na2S
+ NaSH
+ PQ
+ PQ
Na$_2$S
+ PQ
NaSH
+ cysteine
+ PQ,
cysteine
+ PQ
+ Tris-HCl
+ cystine
+ PQ
cystine
+ PQ
Tris-HCl
+ PQ
H$_2$S production
Absorbance at 405 nm
****
****
****
0
50
200
Positive
Ctrl
μM PQ
WT
ΔmetY
ΔmetY, PQ
WT, PQ

Figure S12
